# Nuclear poly(A) tail size is regulated by Cnot1 during the serum response

**DOI:** 10.1101/773432

**Authors:** Richa Singhania, Graeme J. Thorn, Kathryn Williams, Raj D. Gandhi, Clara Daher, Adeline Barthet-Barateig, Hannah N. Parker, Wahyu Utami, Mustafa Al-Siraj, David A. Barrett, Jonathan A.D. Wattis, Cornelia H. de Moor

## Abstract

The poly(A) tail removal from mRNAs introduces a delay between mRNA synthesis and decay. We measured levels and poly(A) tail sizes of serum-induced mRNAs and used mathematical modelling to compare their deadenylation time with the delay in decay and found that they are indeed correlated. Discrepancies between our data and the polyadenylation models at later time points after the peak of induction led us to investigate the size of the poly(A) tails on newly made mRNA. Surprisingly, new serum-induced mRNAs synthesised late in induction had short poly(A) tails (around A_25_) in the nucleus. In addition, newly made constitutive mRNAs had medium sized poly(A) tails (around A_50_). To see if deadenylation was responsible for the new short poly(A) tails, we depleted Cnot1, a subunit of the CCR4/NOT deadenylase. Cnot1 depletion led to slower deadenylation of cytoplasmic mRNAs, as expected, but also decreased transcription and led to longer nuclear mRNA poly(A) tails. These observations implicate CCR4/NOT in regulating both the transcription and the nuclear poly(A) tail size of serum-induced mRNAs. Detection of some chromatin-associated mRNAs with long poly(A) tails suggested that nuclear deadenylation is an early event. Our data show that initial poly(A) tail size of mRNAs can be regulated and is not always 200-250 nucleotides, adding a novel layer to the control of gene expression.

## INTRODUCTION

The stability of an mRNA is a major determinant in the regulation of gene expression. mRNAs encoding proteins involved in the regulation of cellular functions, such as those contributing to gene-specific transcriptional regulation, signal transduction and cell cycle control, are generally short lived. This property makes it possible to rapidly reduce the levels of these mRNAs by shutting down their transcription, as unstable mRNAs disappear promptly in such a situation. In contrast, most mRNAs involved in maintenance functions, such as the general factors required for basal transcription, translation, energy metabolism and protein degradation are quite stable, making them impervious to short term down regulation [1–5]. Presumably, this arrangement allows the cell to save energy by not degrading mRNAs it will certainly need in the future, while allowing for the flexible regulation of other mRNAs as required.

Immediately after transcription, a 200-250 nucleotide polyadenosine (poly(A)) tail is thought to be added to the 3’ end of most mRNAs in the nucleus, with only histone mRNAs escaping the polyadenylation process [6]. In a small number of cases it has been reported that the initial poly(A) tail size is either longer or shorter than this standard size [7–11], but these have heretofore been regarded as exceptions to a near universal rule. In the cytoplasm the poly(A) tail is removed by the activity of two complexes of deadenylating enzymes, Pan2/Pan3 and Ccr4/Not [12]. Removal of the poly(A) tail is a prerequisite for mRNA decay. After shortening of the poly(A) tail mRNAs become a substrate for oligouridylation [13]. This is followed by removal of the 5’ cap (decapping) and rapid decay by the 5’ exonuclease Xrn1 and/or the cytoplasmic exosome [14].

Regulation of the rate of deadenylation is the major factor in determining the stability of several mRNAs, for instance those controlled by the binding of Zfp36/Tristetraprolin to AU rich elements in their 3’UTRs [15]. However, many mRNAs accumulate with relatively short poly(A) tails, suggesting that the decay rate after initial deadenylation also can be modified [16–19]. Indeed, the ELAV-like proteins have been reported to stabilize AU-rich mRNAs after they have been deadenylated [20].

Applying mathematical principles to the still largely descriptive biochemical models is now becoming possible as experimental methods are improving quantitatively. Using a quantitative rather than a descriptive model enables testing of the relative importance of processes that are biochemically linked and to use biochemically correct computational approaches for modelling more complex biological systems, such as gene expression networks. Mathematical models can expose counter-intuitive consequences of biochemical processes or demonstrate that factors are missing from the known biochemical pathways, generating hypotheses which can be tested experimentally [21–23]. As it is not feasible to model every molecular interaction in an organism separately, a key aspect of useful mathematical models is that they include sufficient detail to correctly describe the process and make predictions, but require as few parameters as possible to reduce the types and amounts of data required for analysis.

The requirement for mRNA deadenylation before decay has been known for over 25 years [24, 25] and it was shown in 1993 that there is indeed a delay in the degradation of newly made mRNA in yeast [26]. A detailed mathematical model was developed to represent all the then known steps of mRNA decay in 2001 [27]. Despite this fact, mRNA decay is almost invariably modelled as exponential decay, in which the decay rate is independent of the age of the mRNA, even when rapid changes in gene expression were being studied at a relatively high time resolution [1, 28, 29]. In these models, the fraction of mRNAs that decays per minute (*λ*, the decay rate) is directly related to the half-life and average lifetime of an mRNA (half-life = (ln 2)/*λ*, average lifetime = 1/*λ*) and changes in decay rate are assumed to affect all mRNAs equally, regardless of age. It has been noted previously that mRNA decay kinetics often do not conform to immediate decay and that mRNA age probably plays a role in decay profiles [30].

To improve the existing models, we developed a mathematical model that reflects the biochemistry of mRNA decay but does not require the fitting of too many parameters. To test it, we used a similar biological system to the one in which the role of the poly(A) tail in mRNA stability was discovered: the induction of the immediate early genes with serum in fibroblasts [24, 25]. We obtained data with a high enough time resolution to detect possible delays in decay by fitting a delayed decay model that is a simplified version of a previous model [27], but requires a reduced data set for modelling. We derived the deadenylation time for these mRNAs using a PCR based poly(A) test and a poly(A) tail model. We found the deadenylation time correlated well with the modelled delay in decay and varied between different mRNAs, as expected. However, the poly(A) tail model diverged from the data at the later time points, which led us to question the long standing assumption that all mRNAs are produced with similarly long poly(A) tails. We found that at the peak of transcription, serum response mRNAs were appended with long 150-220 nucleotide poly(A) tails, but when their transcription rates came down, mRNAs with very short poly(A) tails were exported. In addition, we found that all five constitutively expressed mRNAs we examined received much shorter nuclear poly(A) tails than the expected 200-250 nucleotide tails and some also exhibited regulation of their poly(A) tails during induction. To determine if these short poly(A) tails are due to deadenylation, we knocked down the deadenylase subunit Cnot1 and showed that this protein contributes to the regulation of nuclear poly(A) tail sizes as well as to the transcriptional induction of serum response mRNAs. For most genes examined, mRNAs with long poly(A) tails were detected in chromatin associated RNA, suggesting that nuclear deadenylation of mRNAs occurs very soon after transcription. These data indicate that nuclear poly(A) tail size varies and is regulated, providing another mechanism for controlling delayed decay, in addition to the regulation of the cytoplasmic deadenylation rate.

## RESULTS

### Dynamic modelling of mRNA levels

To model the role of the poly(A) tail in gene expression, we chose to study the serum response in NIH 3T3 fibroblasts, because it is a well characterised system with rapid and substantial changes in gene expression [24, 25]. The cells were serum starved for 24 hours and stimulated for 110 minutes, with samples taken every 5 minutes. Two of these time courses were obtained and analysed. To estimate the production of mRNA, mRNA precursor levels were measured (unspliced or intron) and the expression peak of the precursor was assumed to be the result of a variable transcription rate and a constant splicing rate.

We tested 2 models of mRNA decay on these data:

The *immediate decay model (ID)* follows classical exponential decay. In this model pre-mRNAs (*p*) are spliced with rate α to become mRNAs (*m*). These newly made mRNAs are then immediately available for decay with a decay rate λ. The average lifetime of an mRNA in this model is 1/λ:

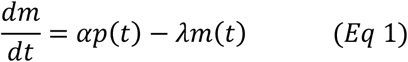

In the *delayed decay model (DD)* all mRNAs are protected from decay for a fixed time τ after their splicing with rate α, becoming susceptible to decay after this period has passed. This model assumes that all the steps that significantly delay decay, which could include export, deadenylation and uridylation, constitute a large series of catalytic steps with similar rates (see Figure 1 for a depiction of this model). Although each of the separate steps will have somewhat different durations for individual mRNA molecules, due to the stochastic nature of enzymatic reactions, the large number of these steps leads to an average protected time τ that varies little between individual mRNAs. Only after this time the mRNA becomes available for a rapid decay process which is initiated by a single reaction, for instance decapping, and therefore behaves as an ordinary enzymatic reaction, leading to exponential decay. By employing this approximation, we arrive at a model in which there are two pools of mRNA, a protected pool m^P^, which is constituted of the mRNAs synthesised during the preceding time τ, and a susceptible pool m^S^ in which the mRNA synthesised before time τ is subject to exponential decay with a rate *λ*. In this model the average lifetime of an mRNA is τ + 1/λ:

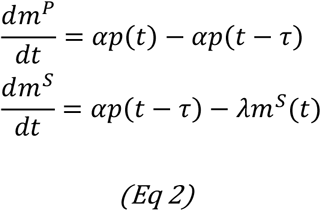

**Figure 1.**
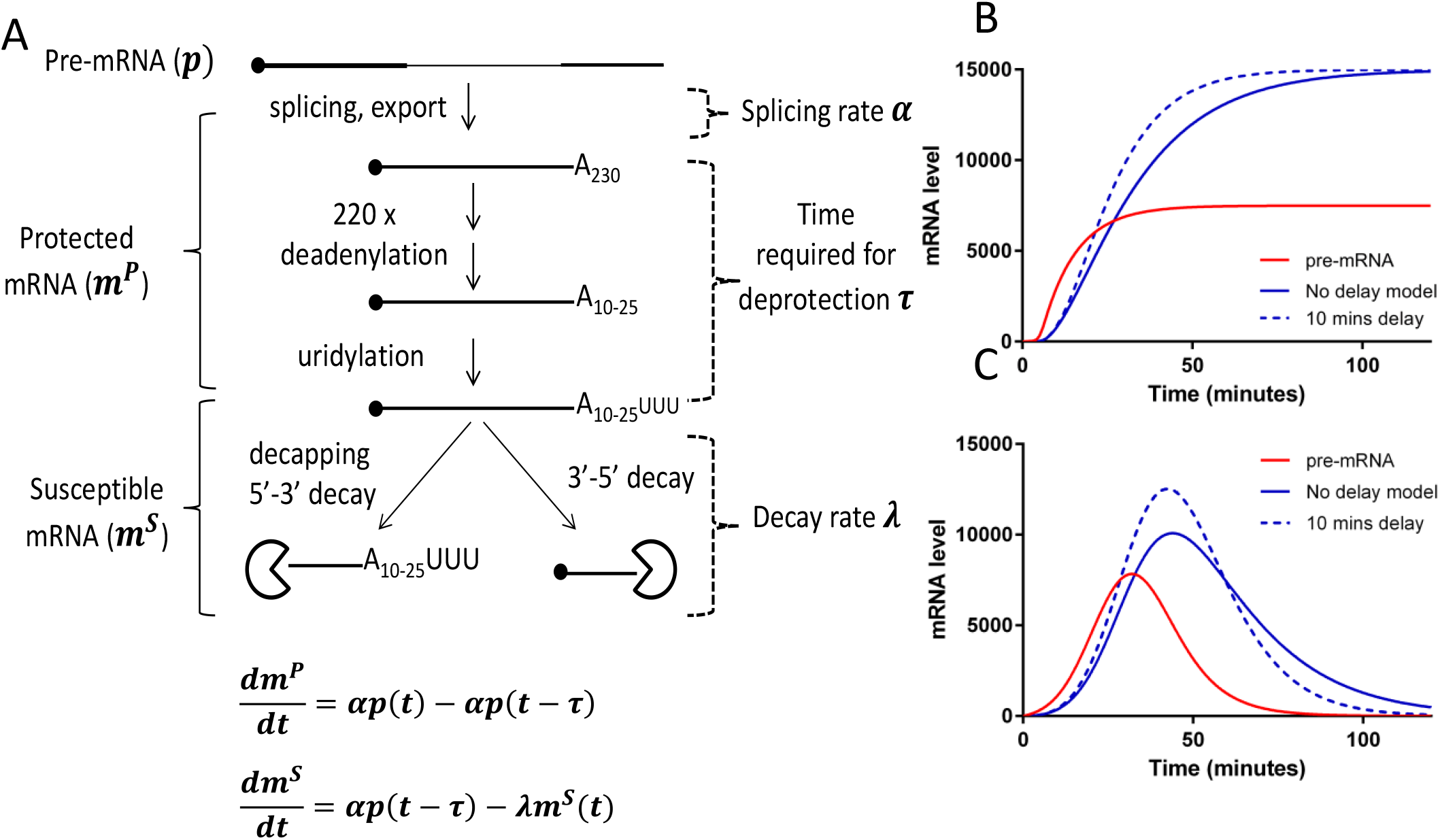
A model for delayed decay of mRNAs. A. A schematic representation of mRNA decay showing how the biochemistry of mRNA decay can be summarised in a pair of ordinary differential equations. B. Simulation of the effect of delayed decay on transcriptional induction. Both mRNAs have a 20-minute lifetime, making the delay 50% of the lifetime. The splicing rate was fixed to be identical. C. Simulation of the same two hypothetical mRNAs being produced from a transient pulse of transcription.

We performed simulations with these models using a hypothetical induction of a pre-mRNA (red curve in Figure 1B). A delay equal to half the average lifetime reduces the time required to reach the steady state (dashed blue line) compared to an mRNA decaying by ID (solid blue line). A similar phenomenon is observed when transcription is switched off in the steady state, with DD leading to more rapid reduction of mRNA levels than ID (not shown). This is because in DD the decay rate for mRNAs that have aged beyond the delay time has to be higher to maintain the average lifetime. A transient pulse of pre-mRNA combines both these effects, leading to a narrower and higher peak of mRNA transcription, as long as the delay time is no longer than the transcription pulse (Figure 1C). These simulations indicate that delayed decay could be particularly important for rapid changes in gene expression, and less so for constitutively expressed mRNAs.

To obtain data for modelling, we first performed quantitative PCR (qPCR) for reverse transcribed spliced and unspliced transcripts of the serum response genes *Fos* and *Fosb*, using standards to obtain absolute numbers. Figure 2 shows the fitted curves for the two time courses for Fos and Fosb. For both models the pre-mRNA input function *p*(*t*) was directly derived from the data for unspliced using a modified Gaussian process fitting procedure (see methods). The optimal values for the parameters (α, *λ*, and, for DD, τ) to fit the mRNA data were derived by performing a random-restart optimisation 200 times and choosing the best fits for use in the figures and table. Because the splicing rate as well as the decay rate can vary when performing this fitting, there is a much more modest difference between the two models than in Figure 1C, where the splicing rate and the average lifetime were kept constant.

**Figure 2.**
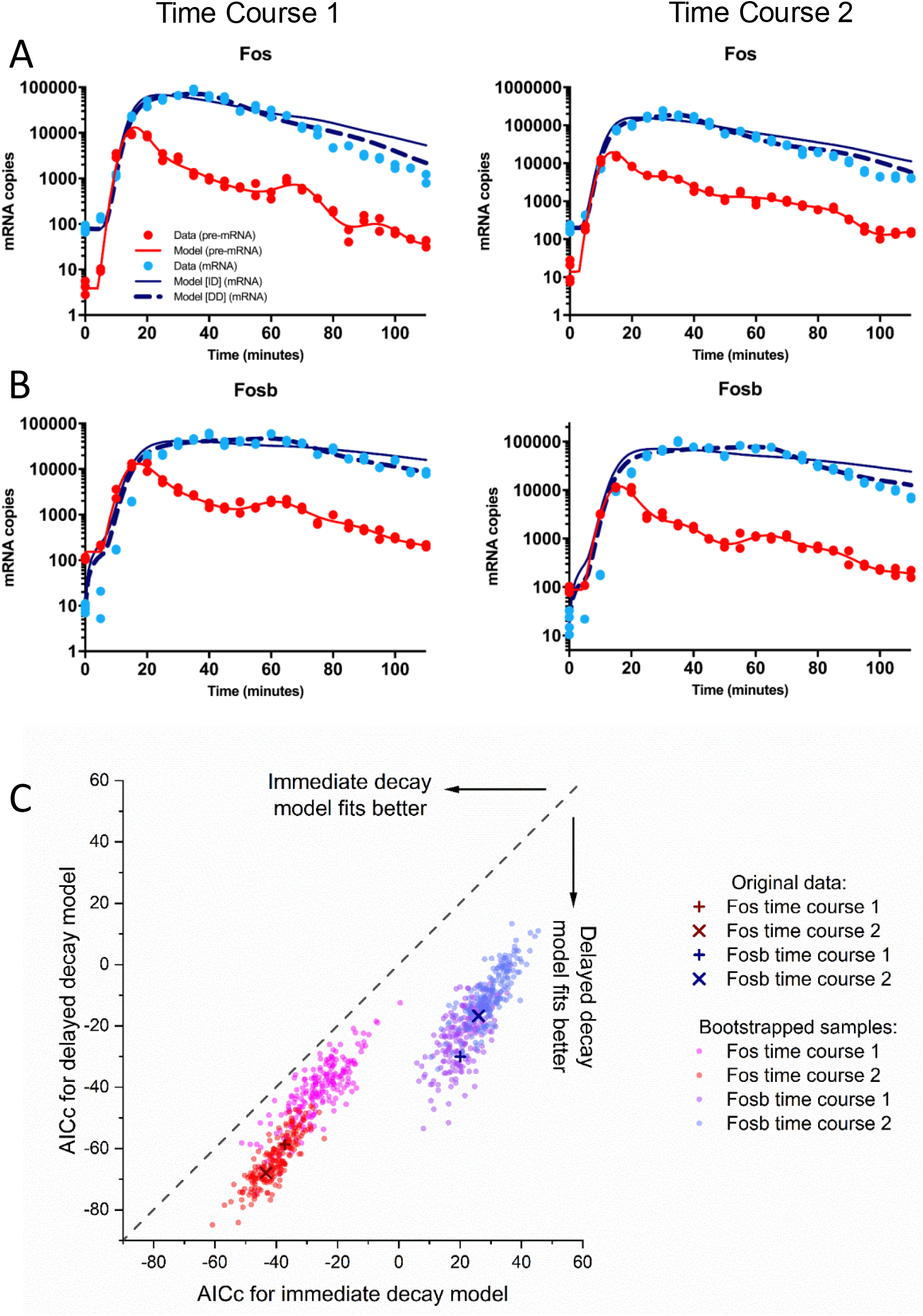
Delayed decay is a better fit for the induction of serum response mRNAs. NIH3T3 cells were serum starved and induced with serum in independent duplicate experiments (Time Course 1 and Time Course 2). RNA was isolated at the indicated times and mRNA and pre-mRNA levels for Fos (A) and Fosb (B) were determined by RT-qPCR using DNA standards. The continuous blue line shows the best fit for immediate decay, the dashed line for delayed decay. (C) Bootstrapped datasets were generated and corrected Aikaike Information Criteria calculated for each bootstrapped dataset. Like the original datasets (+ and ×), all bootstrapped datasets fitted better with delayed decay.

To determine if this difference between the fits of the two models is significant, the fitting was repeated on 200 bootstrapped datasets, which were derived from each original dataset by using the observed variation [31]. This allowed us to test if the improved fit with the DD model justified the introduction of the extra parameter (τ), by calculating the corrected Akaike Information Criterion (AICc) for each model and comparing the AICc for both models for each bootstrapped sample [32]. For all four expression profiles, both the original and all 200 bootstrapped datasets gave a significantly better fit for the DD model. This indicates that the DD model better explains the data (Figure 2C and Table 1). The delay for Fos mRNA is on average 23.5 minutes and for Fosb mRNA 48.5 minutes, demonstrating that the delay time is specific for each mRNA species (Table 2).

**Table 1.** Delayed decay is a significantly better fit for the gene expression data. Corrected Aikaike Information Criteria (AICc) were calculated for the best fits for immediate and delayed decay models for each time course. The relative likelihood indicates how manifold the delay model is more likely to be the correct model vs the immediate decay model. It is calculated as: e^(-(AICc(Direct Decay)-AICc(Immediate Decay))/2)^ [32].

|  | Fos |  | Fosb |  |
| --- | --- | --- | --- | --- |
|  | Time Course 1 | Time Course 2 | Time Course 1 | Time Course 2 |
| AICc Immediate decay | -37.2 | -43.3 | 20.0 | 26.0 |
| AICc Delayed decay | -58.7 | -67.9 | -30.0 | -16.7 |
| Relative likelihood | 4.7E+04 | 2.2E+05 | 7.0E+10 | 1.9E+09 |

**Table 2.** Parameters derived from the best fit models on the quantitative PCR data. The table shows the best fit parameters for immediate and delayed decay, the average lifetime and the total number of transcripts that is made to generate the pulse according to each model.

|  | Immediate decay |  |  | Delayed decay |  |  |  |
| --- | --- | --- | --- | --- | --- | --- | --- |
| | Splice rate $\alpha$ (1/minute) | Average Lifetime $1/\lambda$ (minutes) | Total number of transcripts | Splice rate $\alpha$ (1/minute) | Average lifetime $1/\lambda + \tau$ (minutes) | Delay $\tau$ (minutes) | Total number of transcripts |
| Fos STC1 | 0.8 | 23 | 1.4E+05 | 0.50 | 32 | 22 | 8.9E+04 |
| Fos STC2 | 1.3 | 18 | 4.1E+05 | 0.81 | 27 | 25 | 2.5E+05 |
| Fosb STC1 | 0.3 | 43 | 8.3E+04 | 0.22 | 53 | 47 | 5.8E+04 |
| Fosb STC2 | 0.7 | 41 | 1.4E+05 | 0.44 | 54 | 50 | 9.3E+04 |

The average lifetime of the Fos and Fosb mRNAs was around 10 minutes longer when determined using delayed decay rather than with immediate decay and the splice rates were higher for immediate decay (Table 2). Because the level of unspliced mRNA, which is used as the input function, is the result of the transcription rate and the splicing rate, an increased splicing rate requires increased transcription to maintain the profile. Using the models and the optimised parameters, we found that the number of mRNAs that had to be synthesised to accommodate the models was indeed reduced when delayed decay was used (Table 2).

To obtain information about the decay kinetics for a larger number of mRNAs we applied NCounter analysis to the same two sample sets. This method relies on single molecule detection using RNA specific colour-coded probes and has been reported to be highly sensitive and have a 500-fold linear dynamic range [33]. This allowed us to determine exon and intron counts for 25 serum response genes. To map these data onto our model, we assumed that introns spliced out of the pre-mRNA degraded quickly, thus the intron data generated this way equalled the pre-mRNA levels. This is indeed true when we compare the NCounter intron data and the unspliced qPCR data for Fos and Fosb (Supplementary Figure 1). As both pre-mRNA and mRNA contains exons, the exon data was treated as being equal to the sum of the pre-mRNA and mRNA levels. Data and best fit models for nine examples are shown in Figure 3A and graphs for all profiles are in Supplementary Figure 2. It was clear that despite the expert design by Nanostring, the efficiency of binding of the NCounter probe sets varied widely, e.g. the intron counts for Klf9 were much higher than the exon counts, necessitating the introduction of a scaling factor, which was assumed to be probe specific but identical for all samples. While some expression profiles better follow the data with a DD model (e.g. Cyr61, Egr1, Egr2, Fosb), others converge on short delays, causing the DD and ID curves to overlap (e.g. Id1, Myc). The models yield a variety of delays, which are generally well correlated between the two time courses (Figure 3B). When outliers with poor replication between the two time courses are removed, the delays vary from 0 to 50 minutes. In addition, the decay rates were well correlated between the two models and varied considerably over at least a 10 fold range, indicating that this parameter is also specific for each mRNA, regardless of the model used (Supplementary Figure 3).

**Figure 3.**
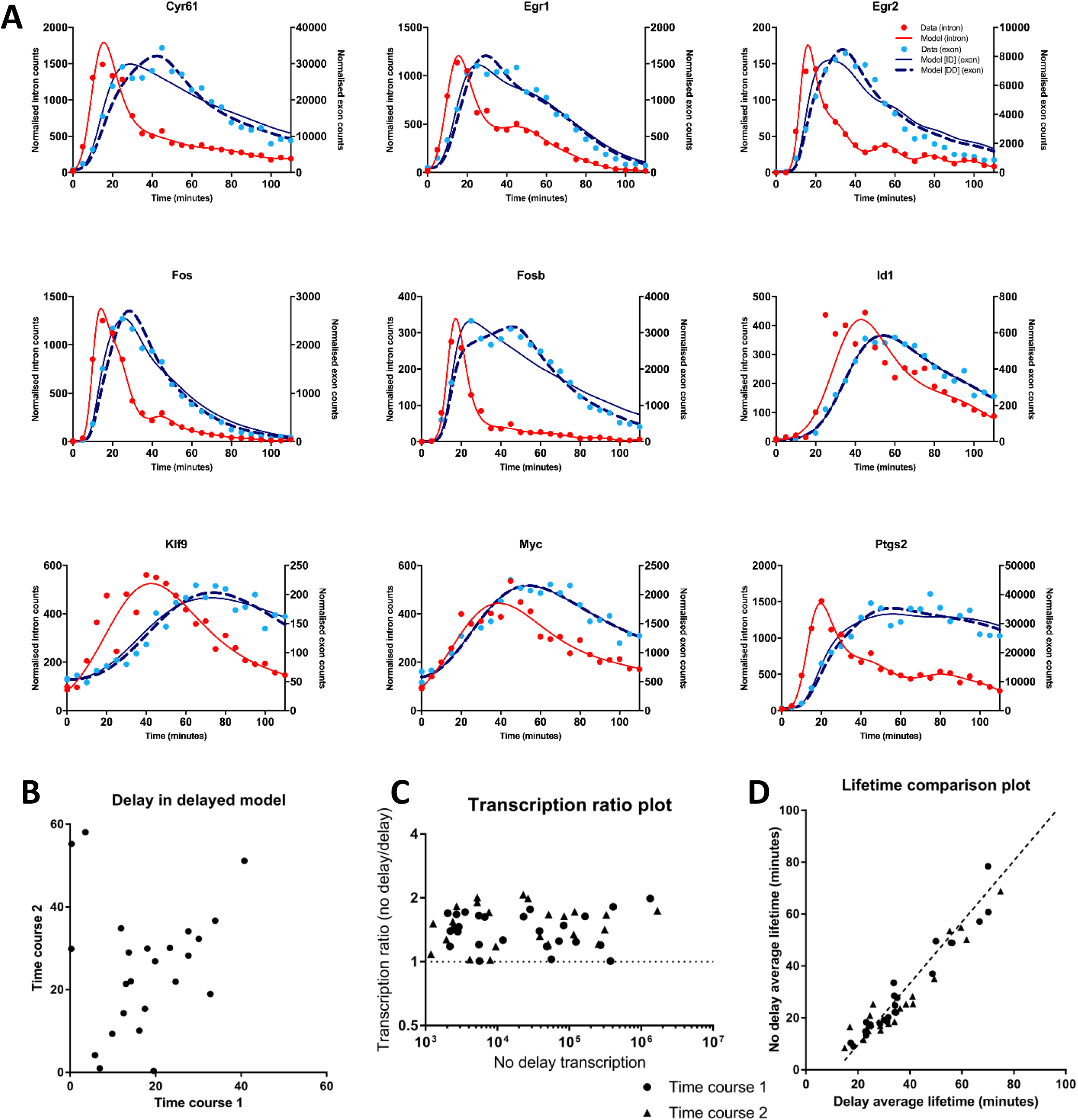
Fitting immediate and delayed decay models to more serum response expression profiles. The RNA samples described in Fig 2 were analysed with NCounter exon and intron probes for 25 serum response genes. A. Examples of fittings of immediate and delayed decay profiles to data for serum stimulation time course 2. B. The best fit delays for each mRNA were plotted against each other for the two time courses. C. The number of pre-mRNA transcripts required to be transcribed (transcription) was calculated for the optimal fit parameters for both models. Transcription for immediate decay (no delay) was plotted vs the ratio of transcription for immediate and delayed decay. D. The average lifetimes for the optimal fitted parameters for delayed and immediate decay were plotted against each other. Duplicates and additional data can be found in Supplementary Figure 2 and 3.

Delayed decay gave marked reductions in the total number of transcripts predicted to be required for the serum response, up to 2-fold fewer transcripts were needed to produce the mature mRNA profile (Figure 3C). Thus, delayed decay can expect to reduce the energy expenditure for the induction of transcription. This was even true in most of the cases where the fitted delay was small and the fitted lines did not appear different for the two models. For instance, there is still a 15% reduction in the number of transcripts required to generate the Myc expression profile shown in Figure 3A, resulting from a 4 minute delay. Logically, a delay would be most advantageous for the genes with the highest transcription level, but no clear correlation between total number of transcripts and delay-mediated savings was observed (Figure 3C).

The estimated average lifetimes varied between 9 minutes for Dusp1 with the ID model to 11 hours for Ereg in time course 2 (with delayed decay). However, given the timespan studied, lifetimes of well over an hour are unlikely to be reliable using these data. There was a good correlation between the average lifetimes for the ID and DD models within this range, with delayed decay generally giving an approximately 10 minute longer lifetime (Figure 3D). The fraction of the lifetime that was occupied by the delay in the DD models was on average 56% but ranged as high as 80% for Btg2. As in the qPCR time courses, the delay for Fos was approximately half that for Fosb, but the absolute numbers differed considerably between the two methods. NCounter data modelling gave an average of 13 minutes for Fos, 31 minutes for Fosb (vs 23.5 and 48.5 minutes for qPCR data). Parameters for all modelled mRNAs can be found in Supplementary Table 1 and further correlations of parameters in Supplementary Figure 3.

### Poly(A) tail distribution modelling

If our model correctly reflects the biochemistry, then the fitted delay time should include the time it takes to deadenylate the mRNAs. This time is determined by the rate of deadenylation and the initial length of the poly(A) tail. To measure the deadenylation time directly, we performed improved poly(A) tests (RL2-PATs, see methods) on the RNA from time courses using gene specific primers for Fos, Fosb, Egr1, Egr2, Id1 and Ptgs2. As shown in Figure 4A, mRNAs with long poly(A) tails appeared for all these mRNAs soon after the induction of transcription and the size of these poly(A) tails gradually declined over time, reaching a minimum size. Visual inspection of when the long poly(A) tails appear and disappear led to estimates of deadenylation times of 15-25 minutes for Egr1, Egr2 and Fos and 30-40 minutes for Fosb and Ptgs2. No clear estimate could be made for Id1 as mRNA with short poly(A) tails is present in all fractions, so the first appearance of deadenylated mRNA cannot be determined. However, the disappearance of the long products suggests that the deadenylation time of Id1 mRNA is shorter than for Fos. Curiously, when PAT products were detected before induction, they appeared to predominantly represent mRNAs with very short poly(A) tails, instead of the mixed distribution of long and short tails that would be expected at the steady state. However, the very low levels of mRNA at these time points made verification of this finding difficult.

**Figure 4.**
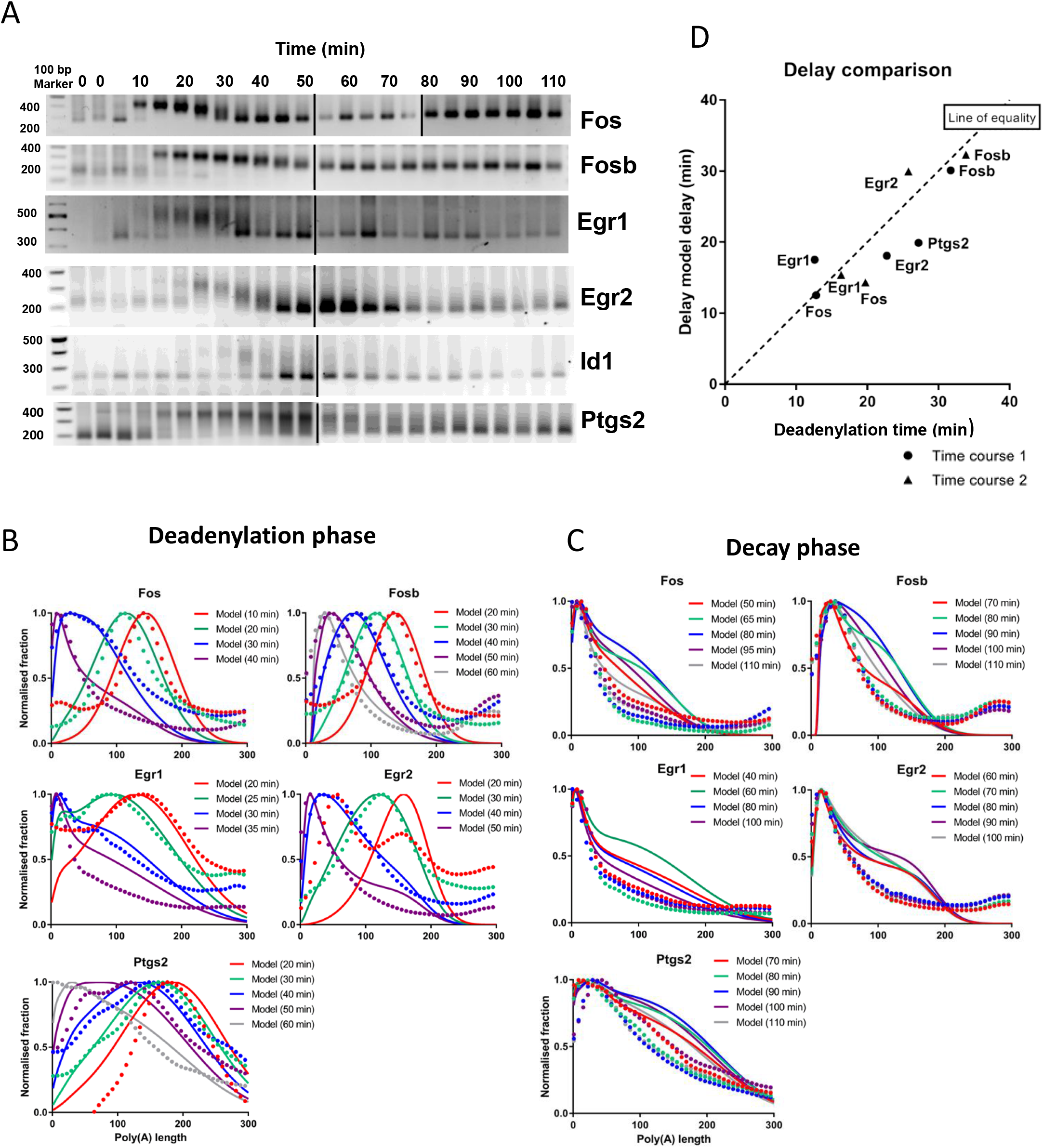
Determination of deadenylation times. A. RNA ligase 2 mediated polyadenylation tests (RL2-PATs) were conducted for 6 mRNAs on the time course 2 samples described in Figure 2 and 3. B, C. The gels were scanned and the data analysed with our poly (A) tail model. All timepoints were used in modelling, but the fits for key times in the deadenylation of each mRNA is shown. D. The deadenylation times from the poly(A) tail model are compared to the delay times from fitting the NCounter data. Duplicate data can be found in Supplementary Figure 4.

To determine the time it takes to deadenylate these mRNAs more accurately, we developed a mathematical model for poly(A) tail size. In this model we use the intron counts from the NCounter data to estimate the production of newly made mRNA and assume that these new mRNAs are produced with a poly(A) tail size distribution that is independent of time. mRNAs are deadenylated at maximum rate *k*^*A*^ for long tails, which is reduced at smaller poly(A) sizes, reflecting a decrease in occupancy of the deadenylase on shorter tails. On average the size of the poly(A) tail at which the deadenylase drops off is *μ*^*A*^, but for individual mRNAs this has a standard deviation of σ^A^. This means that at poly(A) tail size *μ*^*A*^ − *2σ*^*A*^, 97.7% of all mRNAs will have reduced deadenylation rates and the deadenylation of the total population will have significantly slowed. No provision was made in the model for different deadenylase enzymes mediating increased rates of deadenylation at shorter tail sizes [34, 35]. Thus, the deadenylation rate *k*^*A*^ at poly(A) tail size *i* is:

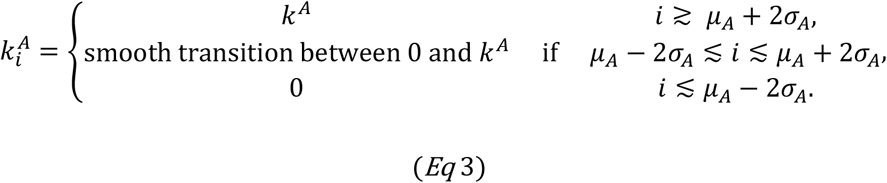

The model also assumes the decay enzymes bind mRNAs at a specific poly(A) tail size with a mean (*μ*^*C*^, at which the average mRNA decays) and a standard deviation *σ*^*C*^. At poly(A) tail size *μ*^*C*^+ *σ*^*C*^, decay can be said to have started. The decay rate *k*^*C*^ at poly(A) tail size *i* is:

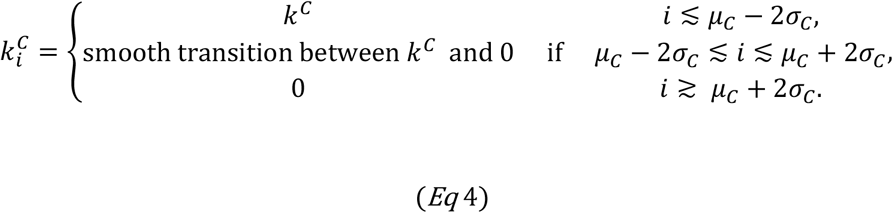

Details of the poly(A) tail model can be found in the Methods.

Selected scans are shown with the fitted plots in Figure 4B and 4C. Early poly(A) tail distributions (in red in Figure 4B) fitted less well because of poor signal to noise ratios or remaining signal from before induction (Egr2), but still matched the peak sizes. The later fits largely captured the poly(A) tail size changes in the data, evident from the movement of the band during the deadenylation phase. It is apparent from the figures that the band moved more slowly for Fosb and Ptgs2 compared to the Fos, Egr1 and Egr2 results, which corresponded with the movement of the longer tail band in the gel images. In contrast, consistent discrepancies were seen in all the fits in the decay phase (Figure 4C), after the mRNA generated during peak expression had been deadenylated. Less polyadenylated mRNA transcribed late in the response was detected in the data than predicted by the models.

The parameters obtained for the fits for PAT data for both time courses are shown in Supplementary Table 2. The parameters for Egr1 timecourse 1 are clear outliers, with the exception of the deadenylation time, perhaps because we did not detect the early and late PAT products for this mRNA in this timecourse (see Supplementary Figure 4). Removing this dataset, decay starts at a tail size of 20 (average of μ^C^ + σ^C^ over the 8 remaining datasets), which matches published data on the start of uridylation [17] and correlates with the 25 nt binding length for a single poly(A) binding protein [36, 37].

To determine if deadenylation contributes significantly to the delay, we plotted the deadenylation time from the poly(A) model against the delay from the delayed decay modelling on the NCounter data. As can be seen in Figure 4D, the deadenylation times were on average somewhat longer than the delay times (they are on the right of the line of equality), which may be due to decay starting before deadenylation is complete. The delay and the deadenylation time were very well correlated (R^2^ = 0.74, P=0.0031), clearly indicating that the delays derived from the gene expression modelling at least in part reflect the deadenylation times.

### Nuclear poly(A) tail size is regulated

The poor fit of the poly(A) tail modelling for the late time points (Figure 4C) and the detection of mRNAs with discrete short poly(A) tails before induction (Figure 4A) led us to question the assumption that mRNAs are made with a constant poly(A) tail distribution throughout the serum response. To investigate the possibility that initial poly(A) tail size is shorter during the late phase of the induction, we decided to label cells with 4-thiouridine before lysing the cells and examining intracellular levels of thioUTP to determine optimal labelling conditions (Supplementary Figures 7 and 8). As a result, we decided to label NIH-3T3 cells with 250 μM thiouridine for 10 minutes. The cells were either untreated or treated with serum for 5, 20 or 50 minutes, giving endpoints of untreated (0’), 15 minutes, 30 minutes or 60 minutes serum treatment. The 4-thiouridine RNA was isolated on streptavidin beads after biotinylation (bound fraction) and poly(A) tests were performed. Quantitative PCR for the spliced and unspliced forms of a constitutive expressed mRNA confirmed that newly made RNA is enriched in the bound fraction (Supplementary Figure 5). Our improved poly(A) test allowed us to measure poly(A) tail sizes of individual mRNAs in these minute amounts of RNA for the first time (see Methods). Poly(A) tail sizes for the serum response mRNAs (Egr1, Egr2 and Fos) in total and unbound fractions were similar (Figure 5A). At 30 minutes, the bound fraction had longer poly(A) tails than the total RNA, also confirming that the bound fraction is enriched for new mRNA. However, at 60 minutes the sizes poly(A) tails of bound serum response mRNAs were predominantly short, suggesting that when transcription rates drop, serum response mRNAs are made with short poly(A) tails.

**Figure 5.**
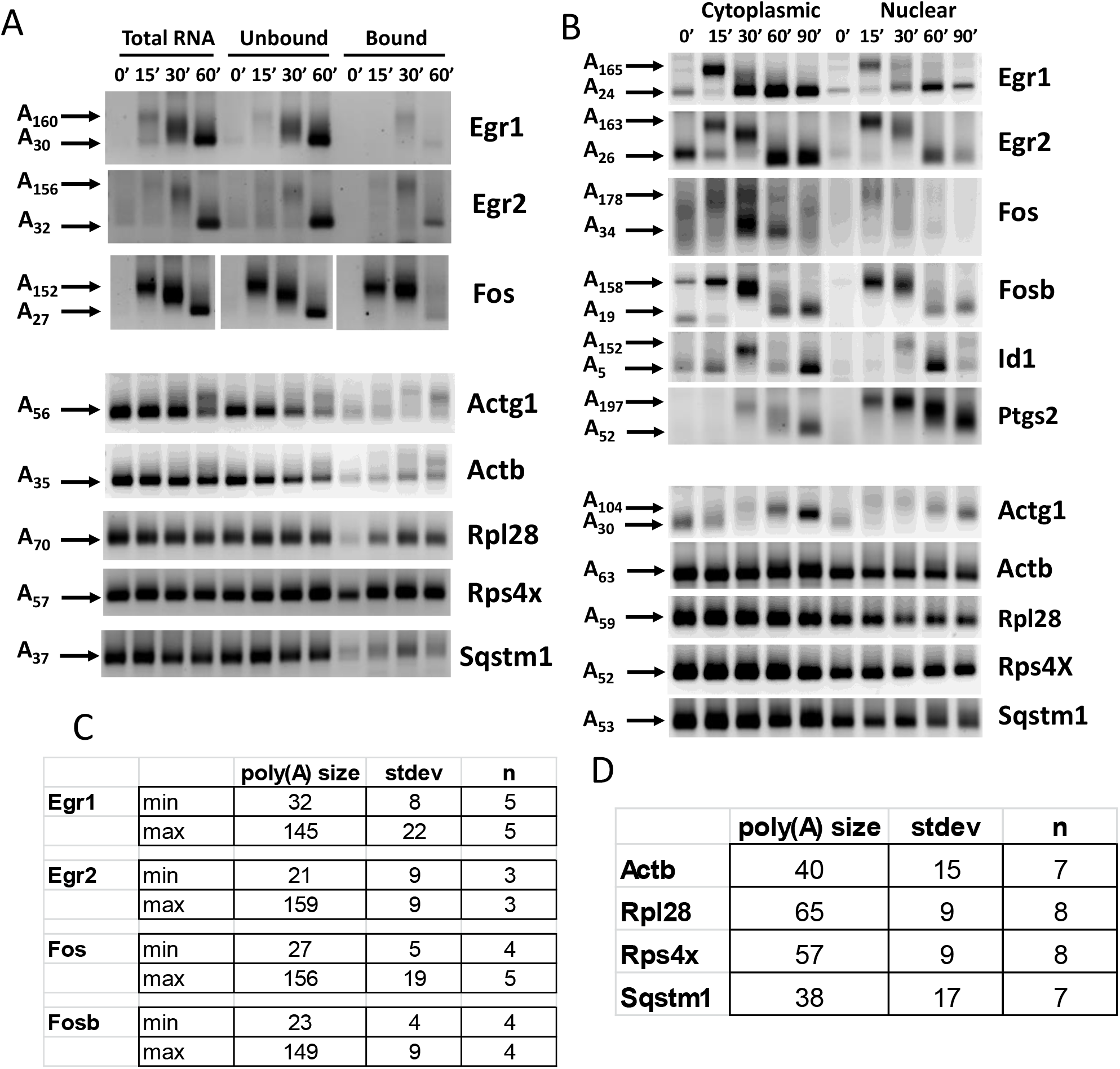
Nuclear poly(A) tail size is regulated. A.NIH3T3 cells were serum starved and incubated with 4-thiouridine for 10 minutes, either without stimulation with serum (0’) or from 5 to 15 minutes (15’,) 20 to 30 minutes (30’) or 50 to 60 minutes (60’). Total RNA was isolated and labelled RNA was isolated from a fraction (bound). RL2-PAT was performed for the indicated mRNAs. Arrows indicate maximum and minimum modal poly(A) tail sizes, determined using quantitative gel scanning as described in the Methods. B. Nuclear and cytoplasmic RNA was isolated from NIH3T3 cells induced with serum for the indicated times and subjected to RL2-PAT. Arrows indicate maximum and minimum modal poly(A) tail sizes as determined using quantitative gel scanning. C. Maximum and minimum poly(A) tail sizes during the serum response for four responding mRNAs. D. poly(A) tail sizes for constitutively expressed mRNAs in uninduced cells. Additional replicates of A and B can be found in Supplementary Fig 6.

We found to our surprise that in the serum deprived cells (0’), at steady state, newly made constitutively expressed mRNAs for Actb, Rpl28, Rps4x and Sqstm1 do not have 200-250 nucleotide poly(A) tails, but instead have very similar size distributions in bound and total RNA (Figure 5A). These distributions have average modal sizes of around 50 nucleotides. The similarity in these distributions between new and total RNA also suggests that these mRNAs are not gradually deadenylated over their lifetimes, as more short products should then be found in the total than in the bound RNA. In addition, the Actg1 and, to a lesser extent, Actb mRNAs underwent clear increases in poly(A) tail size during serum stimulation, with perhaps a small increase for Sqstm1.

To confirm by another method that regulation of initial poly(A) tail size is occurring, we separated nuclear and cytoplasmic RNA. We checked that the fractionation was working as intended by determining the enrichment of unspliced precursor mRNA over spliced mRNA by RT-qPCR (Supplementary Figure 9). Poly(A) tests showed that for Egr1, Egr2, Fosb and Id1 there was a clear shortening of the poly(A) tails in the nucleus after the peak of induction had passed (Figure 5B). For Egr1 and Egr2, short products were detected in the nuclear fractions before induction, indicating that the background transcription of these genes also produces mRNAs with short poly(A) tails. For Fos, we could not detect nuclear mRNA with short tail lengths in this experiment. These data indicate that there is regulation of nuclear poly(A) tail length for serum response mRNAs during their induction.

When we examined the constitutively expressed mRNAs, the poly(A) tails of Actg1 mRNA were again clearly regulated during the serum response, while very little increase in tail length was seen for Actb (Figure 5B). Once more, Sqstm1 appeared to have a small increase in tail size. Size distributions in the untreated samples appeared the same in cytoplasmic, nuclear and total RNA. The modal sizes are significantly shorter than the maximum 150 nucleotide poly(A) tail size found for serum response genes (2-tailed t-Test, p= 10^−18^). However, the average 51 nucleotide poly(A) tail on constitutively expressed mRNAs is still significantly longer than the tails on the serum response mRNAs late in induction, which are on average 26 nucleotides (2-tailed t-test, p=6.9*10^−8^), as summarised in Figure 5C and 5D. The similarity of the distributions in the nuclear and cytoplasmic fractions again indicates that there is no gradual deadenylation of the housekeeping mRNAs.

### Cnot1 regulates transcription and nuclear poly(A) tail size of serum response mRNAs

The regulation of nuclear poly(A) tail size and the transcriptional pulse of serum response genes appeared temporally tightly coupled, suggesting a co-regulation mechanism. As described above, the mammalian CCR4/NOT deadenylation complex regulates cytoplasmic poly(A) tail size. However, it also has reported effects on transcription, making it a good candidate for mediating this regulation [38–41]. We therefore knocked down the mRNA for the scaffold subunit of CCR4/NOT, Cnot1, using an siRNA transfection approach. Remarkably, a reduction in Cnot1 levels caused a substantial reduction in the induction of serum response mRNAs as well as in the steady state levels of ribosomal protein mRNAs, such as Rpl28 (Figure 6A). This appears to be part of a general repression of transcription as unspliced mRNAs of both serum response and housekeeping mRNAs were also reduced (Figure 6B). The timing of the unspliced mRNA production of serum response mRNAs was unaltered, but the mature mRNAs persisted longer, as would be expected when deadenylation is slowed down, increasing the delay in decay. Indeed, mRNAs in Cnot1 knockdown cells had longer poly(A) tails at all time points tested (Figure 6C). To investigate if Cnot1 also affects nuclear poly(A) tail sizes, we performed poly(A) tests on cells stimulated with serum for 60 minutes, the time when the poly(A) tail size difference between control and Cnot1 knockdown cells is greatest. As can be seen in Figure 6D, while the difference in poly(A) tail sizes between the control and the knockdown is clear, poly(A) tails are equally long in nucleus and cytoplasm. These data implicate Cnot1 in the transcription and the regulation of nuclear poly(A) tail size of serum response mRNAs.

**Figure 6.**
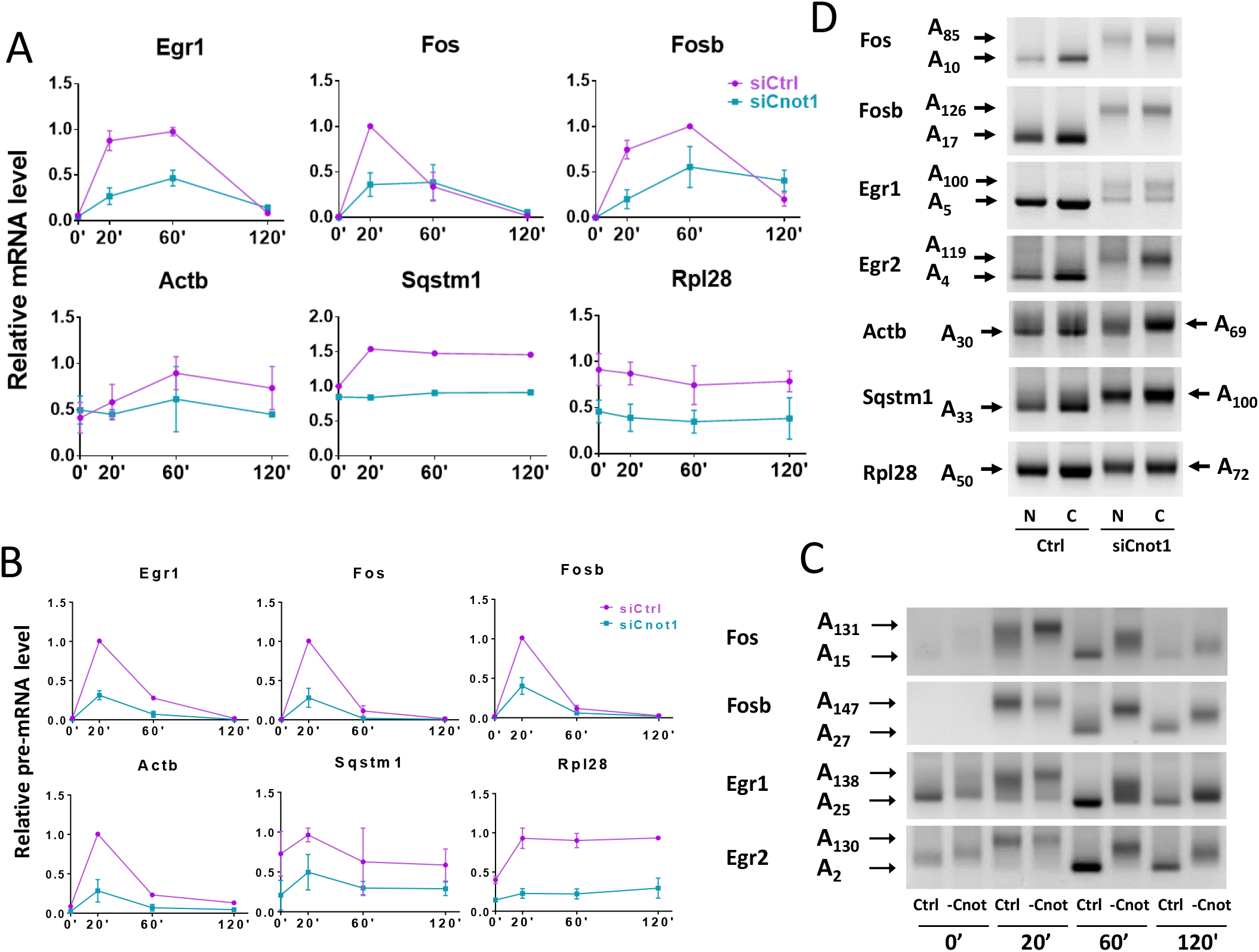
Cnot1 knockdown affects serum response gene transcription and nuclear poly(A) tail size. A. NIH3T3 cells were treated with control siRNAs or siRNAs against Cnot1. Cells were serum starved, either without stimulation with serum (0’) or with serum addition for 20 minutes (20’), 1 hour (60’) or 2 hours (120’). Total RNA was isolated, reverse transcribed and qPCR performed for the indicated mRNAs (A) and pre-mRNAs (B). C. RL2-PAT was performed for the indicated mRNAs. Arrows indicate maximum and minimum modal poly(A) tail sizes, determined using quantitative gel scanning as described in the Methods. D. Nuclear and cytoplasmic RNA was isolated from NIH3T3 cells induced with serum for 60 minutes and subjected to RL2-PAT. Arrows indicate maximum and minimum modal poly(A) tail sizes as determined using quantitative gel scanning. Additional replicates and controls can be found in Supplementary Fig 14.

The most likely mechanism by which Cnot1 affects nuclear poly(A) tail sizes is by stimulating deadenylation of newly made RNA. To investigate if the mRNAs with short poly(A) tails that we detect in the nucleus are generated from long-tailed precursors, we isolated chromatin associated, nucleoplasmic and cytoplasmic RNA and performed RL2-PAT. As can be seen in Figure 7, we could detect long tailed products for most mRNAs in the chromatin fractions, except for Rpl28 and Fos. In most cases, the products with long poly(A) tails were in the minority even in the chromatin associated fraction, with the exception of Sqstm1, for which the poly(A) tail was predominantly long in the chromatin and short in the nucleoplasm. Sanger sequencing confirmed that the long Sqstm1 product is not the product of alternative polyadenylation but genuinely represents Sqstm1 mRNA with a long poly(A) tail. These data suggest that nuclear deadenylation of mRNAs takes place on the chromatin soon after transcription.

**Figure 7.**
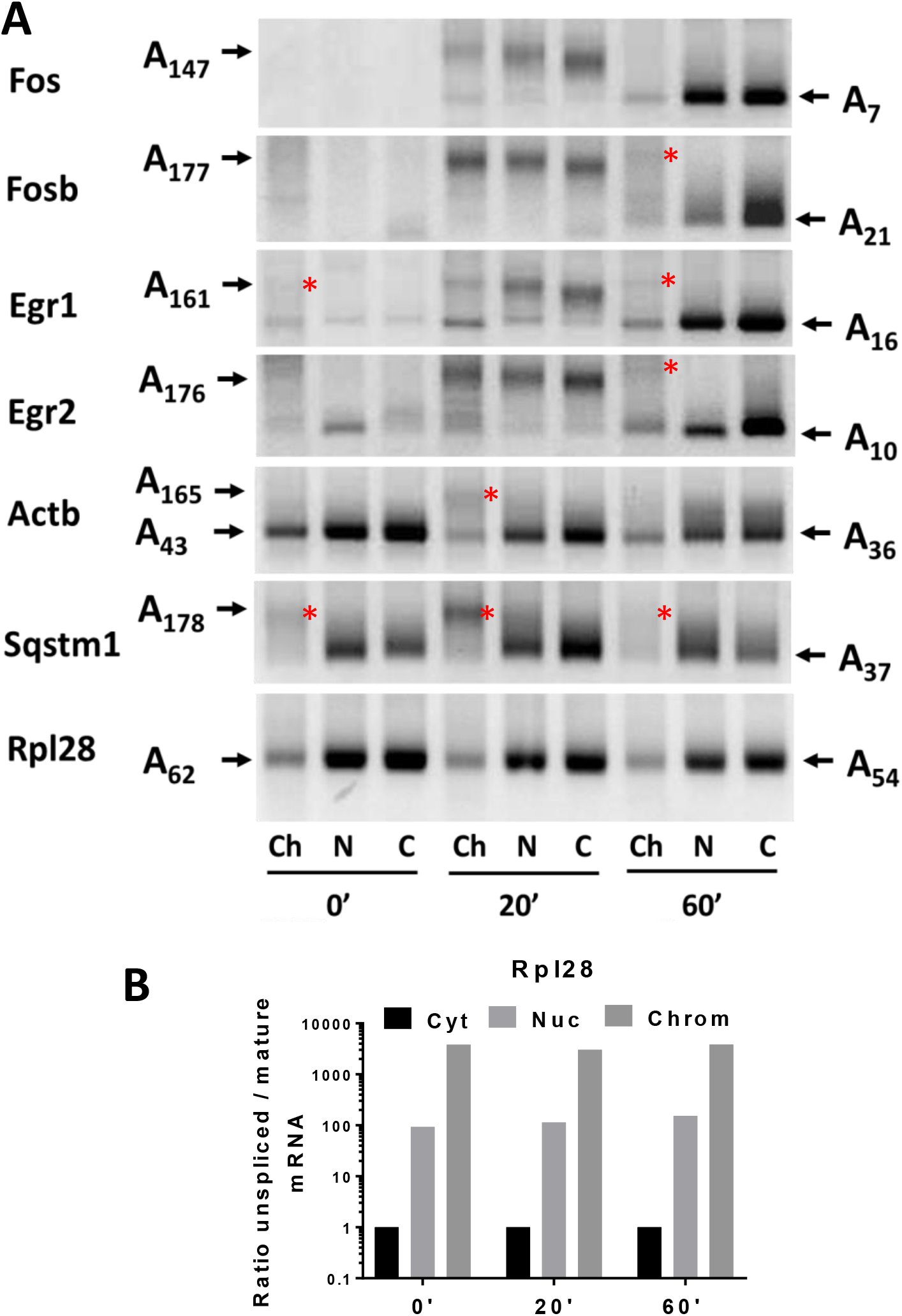
Most mRNAs with short poly(A) tails appear to have long-tailed precursors on the chromatin. A. Serum starved NIH3T3 cells were not treated (0’) or treated with serum for 20’ or 60’. Cells were fractionated into chromatin, nucleoplasm and cytoplasm and RNA was isolated. RL2-PAT was performed on this RNA for the indicated mRNAs. Red stars indicate PCR products which indicate that short-tailed mRNAs have long-tailed precursors. B Enrichment of uspliced Rpl28 mRNA in the chromatin and nucleoplasmic fractions. An additional replicate can be found in Supplementary Fig 15.

## DISCUSSION

In this paper we investigated the role of the poly(A) tail in serum response gene expression using biochemistry and mathematical modelling. Our mathematical versions of classical biochemical models allowed us to identify a discrepancy between the data and the biochemical model. Surprisingly, this led to the observation that the nuclear poly(A) tail of serum response mRNAs is highly regulated during their induction. In contrast, constitutively expressed mRNAs had shorter nuclear poly(A) tails that did not appear to be subject to gradual deadenylation in the cytoplasm. Knockdown of Cnot1 implicated the CCR4/NOT complex in the regulation of nuclear poly(A) tail size, as well as in general transcription.

Only one case of a change in initial poly(A) tail size has been previously reported to our knowledge, and this was associated with gene induction of the eNOS mRNA [10, 11]. The poly(A) tail size regulation of vasopressin mRNA, for which the regulatory mechanism was not investigated, could potentially also be caused by changes in initial poly(A) tail size [42, 43]. Our data indicate that initial poly(A) tail size is regulated for at least 6 mRNAs after serum stimulation (Figure 5A and B), and probably is similarly affected for many more serum response genes. This novel regulatory mechanism may contribute to the previously reported “nuclear imprinting” of mRNAs, in which their cytoplasmic stability is pre-determined during transcription [44]. Indeed, the CCR4/NOT complex has been implicated in such co-regulation in yeast [41]. Our observation that transcription of serum response genes also requires Cnot1 indicates that these genes are regulated by CCR4/NOT and favours a model by which high transcription is coupled to rapid mRNA degradation through this complex.

As expected, Cnot1 knockdown delays the cytoplasmic deadenylation of the bulk of the serum response mRNAs that are produced with long poly(A) tails at the peak of transcriptional induction. However, our data also show that Cnot1 knockdown affects the nuclear poly(A) tail sizes of both off peak serum response and constitutively expressed mRNAs. The fact that we find PAT products indicating long poly(A) tails on the chromatin for most mRNAs indicates that deadenylation occurs very soon after polyadenylation, probably co-transcriptionally. This process has to be extraordinarily fast in comparison to cytoplasmic deadenylation, as the dwell time of mRNAs on the chromatin is much shorter than in the cytoplasm. Known general regulators of nuclear poly(A) tail size are the nuclear poly(A) binding proteins Pabpn1 and Zc3h14 and the non-specific RNA binding protein nucleophosmin (Npm1) [45, 46]. However, we clearly need a means of gene specific regulation. A good candidate for the regulation of the initial poly(A) tails size of serum response mRNAs is Zfp36 (tristetraprolin), a protein that is known to bind the AU rich elements that are common in serum response mRNAs and regulates their stability in the cytoplasm [15]. Zpf36 has also been shown to interact with Pabpn1 and inhibit nuclear polyadenylation [47]. In addition, U rich element binding proteins from the Elavl (Hu) family have been shown to affect the addition of the poly(A) tail *in vitro* [48] and Actb1 mRNA has extensive binding sites for this protein in its 3’ UTR [49].

It is unclear to us why the two different ways of quantitating the same serum response time courses give different delays in decay while there is reasonable correspondence between the two time courses measured with the same technique (Figure 2 and Supplementary Table 1). Variation in amplification efficiency generates systematic logarithmic errors in qPCR data, and is therefore likely to amplify the differences when mRNA levels vary widely, as in our experiments. In contrast, NCounter data are expected to have linear errors only. However, this should have been countered by our use of calibration standards in our qPCR assays. There is an approximately twofold difference between the delays in Fos and Fosb mRNA decay in both methods, so relative values appear to be maintained regardless of the experimental method.

The change in the nuclear poly(A) tail size of mRNAs produced during the course of the serum response could have consequences for the expression profile by producing mRNAs with a shorter delay in decay later in the response. To investigate this, we adapted our delayed decay model to incorporate a change in the delay from long to short after the transcriptional induction has peaked. When we used parameters similar to those derived for Fos and Egr1, the simulated expression profile underwent a very marked sharpening of the expression peak, reducing the trailing of the mRNA regulation behind the transcription regulation (Figure 8). It is very possible that this phenomenon is shortening the fitted delays in the delayed decay model and the fitted deadenylation times in the poly(A) tail model, because these models will mistake rapidly decaying newly made mRNAs with short poly(A) tails for older mRNAs that have been deadenylated. With our current datasets and models, we can therefore not reliably distinguish between delayed decay with a constant delay τ, as described in Figure 1, and delayed decay with a variable delay due to a change in nuclear poly(A) tail size. This is especially true if mRNAs with shorter poly(A) tails start being produced before the majority of polyadenylated mRNAs is fully deadenylated. Nevertheless, the delay parameter τ (Equation 2) is clearly an important indicator of whether delayed decay is facilitating changes in gene expression and using this simple model would add value in all mathematical modelling of rapid changes in gene expression and is likely to lead to more accurate determination of the average lifetimes of mRNAs.

**Figure 8.**
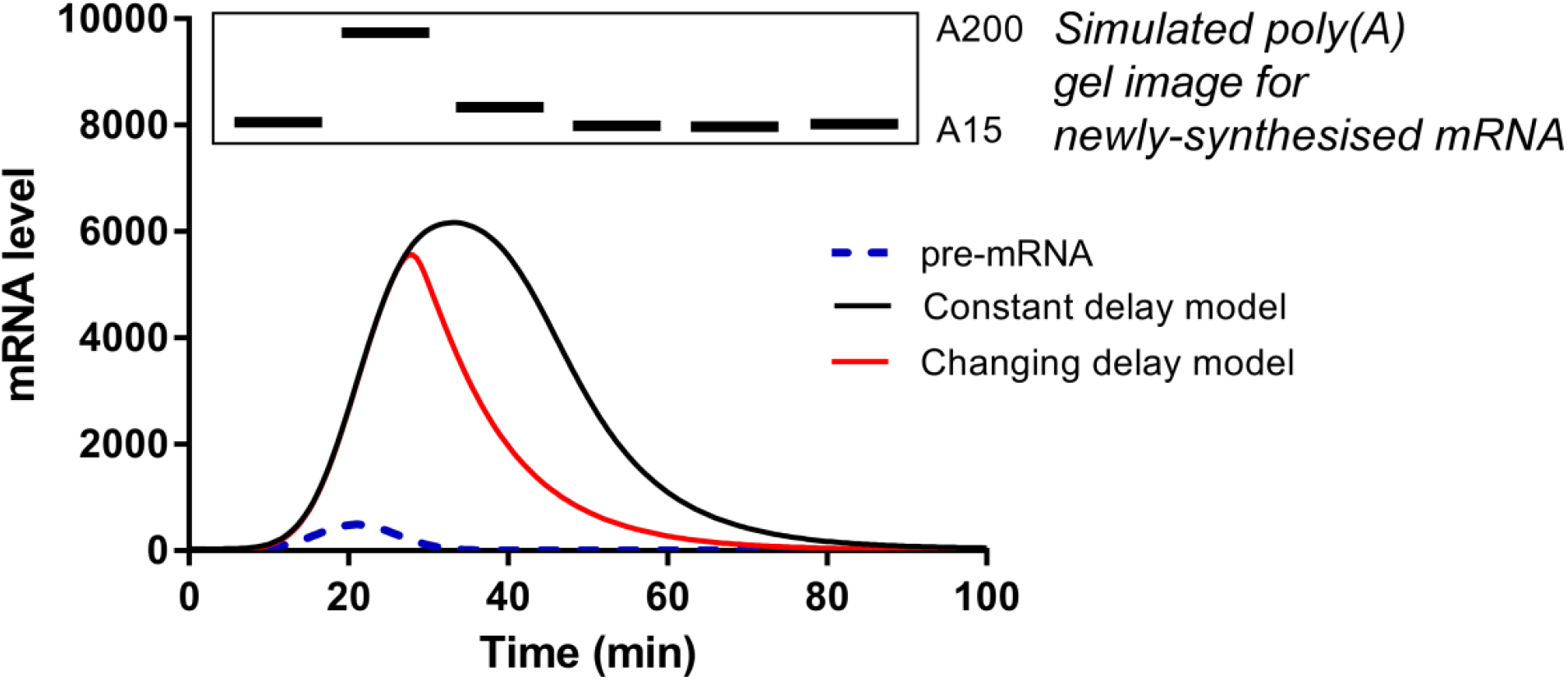
Modelling of the effect of changes in initial poly(A) tail size. The original delayed decay model was compared in a simulation to a model incorporating a change in delayed decay due to the synthesis of mRNAs with short poly(A) tails late in the response.

As shown in Figure 1, mRNAs that do not change rapidly in abundance have no particular need for delayed decay. In addition, the predicted average lifetimes for immediate and delayed decay are similar for longer lifetimes (Figure 3D), and the transcriptional advantage of delayed decay would be lost during constitutive expression. This is because the average lifetime alone determines the transcription required to maintain the steady state, regardless of the decay pathway followed. Remarkably, we find that all constitutively expressed mRNAs we examined in untreated cells did not receive 150-250 nucleotide poly(A) tails, but instead obtain medium sized tails, around 50 nucleotides (Figure 5). The tails on these mRNAs were slightly elongated in the Cnot1 knockdown in both nucleus and cytoplasm but did not exceed 80 nucleotides and did not show differences between nuclear and cytoplasmic mRNA (Figure 6). In addition, the poly(A) tail profiles suggest that these tails are not gradually removed during their lifetimes. Indeed, it has been shown, by us and others, that many stable housekeeping mRNAs accumulate with relatively short poly(A) tails, but this has been attributed to cytoplasmic deadenylation of longer initial poly(A) tails [16–19, 50]. Together, these data suggest that constitutively expressed mRNAs have a distinct poly(A) metabolism from serum response mRNAs and may lack delayed decay.

A few cases of initial poly(A) tail sizes shorter than 150-250 nucleotides have been previously reported [7–9], but these were considered to be the exception rather than the rule. We, however, found nuclear poly(A) tail sizes of less than 70 nucleotides for all 11 mRNAs we examined under at least one condition, indicating that it is probably quite common (Figure 5). The constitutively expressed mRNAs have medium sized nuclear tails of 50 nucleotides in serum starved samples, while the serum response genes have very short 25 nucleotide tails before and/or after induction, indicating that these are differentially regulated. These sizes suggest that the constitutively expressed mRNAs can bind two copies of the cytoplasmic poly(A) binding proteins, while serum response mRNAs generated after the transient induction can only bind one poly(A) binding protein [36, 37]. A high throughput determination of poly(A) tail sizes in nucleus and cytoplasm or on briefly metabolically labelled and total RNA should be able to establish definitively how common short initial poly(A) tails are and how many different classes of initial poly(A) tail size can be distinguished [17, 18, 50].

The Schoenberg lab has identified a poly(A) limiting element (PLE) in the 3’ UTR of several mRNAs with a very short initial poly(A) tail (17 nucleotides) and shown that binding of the splicing factor U2AF to this element promotes the production of long, rather than short, poly(A) tails [7–9]. PLEs are not thought to be common and convey very short oligo(A) tail sizes, so other mechanisms of controlling poly(A) tail size are likely to exist. The variety of initial poly(A) tail sizes and size changes we observe also indicate that there are probably multiple mechanisms for regulating the nuclear poly(A) tail size, expanding the repertoire of gene regulatory mechanisms. We conclude that the regulation of nuclear poly(A) tail size is likely to affect many mRNAs and merits further investigation.

## MATERIALS AND METHODS

### Tissue culture, RNA isolation, RT-qPCR and NCounter analysis

NIH 3T3 fibroblasts were obtained from the ECACC and grown in DMEM with 2 mM glutamine and 10% new born bovine serum as recommended. All experiments were done with a single serum batch, except for Figure 5C. Testing for mycoplasma was performed with the Biotool Quicktest kit v2. For serum induction 10^6^ cells were seeded in 10 cm plates and the medium was changed to 0.5% serum 24 hours later. 22-24 hours after the medium change, the serum response was induced with the addition of 10% new born calf serum. Every 5 minutes, plates were drained of medium and placed on ice. Within 15 minutes of being placed on ice, cells were washed twice with cold PBS, collected by scraping and lysed in the buffer provided by the Nucleospin RNA isolation kit (Mackerey-Nagel) or the Reliaprep miniprep kit (Promega). RNA was isolated according to the manufacturer’s instructions, except that the DNase incubation was lengthened to 1 hour to reduce genomic DNA contamination.

To isolate nuclear and cytoplasmic RNA, a cell pellet containing up to 10 million cells was suspended in 400 μl hypotonic buffer (10 mM HEPES pH7.9, 10 mM KCl, 0.1 mM EGTA, 0.1 mM EDTA, 1 mM DTT, 0.5 mM PMSF). Cells were lysed with 25 μl 10% Igepal. After centrifugation, the pellet was washed three times with hypotonic buffer (nuclear fraction) and the supernatant was precipitated with isopropanol and centrifuged to generate a pellet (cytoplasmic fraction). Both pellets were dissolved in lysis buffer from an RNA isolation kit and RNA was isolated as described above. The nuclear RNA received an additional DNAse treatment. A detailed protocol is available on request. Fractionation of cells into chromatin, nucleoplasm and cytoplasm was performed as described [51]. For RT-PCR, 500 ng of RNA was reverse transcribed with Superscript III (InVitrogen) according to the manufacturer’s instructions.

### Quantitative PCR

Real-time quantitative PCR (qPCR) was performed on a Rotor-Gene Q 2plex HRM Platform (Qiagen) using SYBR Green as the detection dye. PCR amplification was performed in 10 μL reactions comprising of 1X GoTaq^®^ qPCR master mix (Promega), 1 μM of gene specific forward and reverse primers (sequences are provided in Supplementary Table 3) and 2 μL of 1:10 diluted cDNA as template. PCR thermal profile comprised of (1) hold: 95°C for 10 minutes, (2) cycling: 95°C for 10 seconds, 60°C for 15 seconds and 72°C for 20 seconds repeated for 40 cycles (3) melt: 75 −95°C to determine the melting curve. When new primer pairs were introduced, sizes of PCR products were checked on agarose gels in both highest and lowest abundance samples and the efficiency of the amplification verified in a serial dilution of cDNA. When relative mRNA levels were determined, as in the actinomycin D treatments, the ΔΔCt method was used with normalisation to Gapdh.

To obtain standards for absolute quantitation, qPCR products were cloned into pGEM®-T plasmid (Promega) and transformed into TOP10 chemically competent *E. coli* cells (Invitrogen). Plasmid DNA was isolated using PureYield™ plasmid miniprep system (Promega), according to manufacturer’s protocol. To estimate the number of mRNA copies/cell, Ct values of samples were compared with a standard curve generated by dilution series of target plasmid (pGEMT-UnFos, pGEMT-Fos, pGEMT-UnFosb or pGEMT-Fosb) and reference plasmid (pGEMT-Gapdh) using Rotor-gene Q software (Qiagen). Primer sequences are found in Supplementary Table 3.

### NCounter analysis

NCounter probes for an intron and an exon of 45 test genes were designed by Nanostring (Supplementary Table 4). 10 genes were classed as control genes and their exon and intron data used for normalisation. Plk2 was weakly induced and left out of the dataset. Of the remaining 34 genes 6 were expressed below detection levels for either intron or exon counts and were removed from the analysis. One further profile was discarded because the intron levels were approximately constant over time, while the exon levels showed a typical induction profile (Has2). Two serum response gene profiles indicated that there was no precursor-product relationship between the intron and exon counts, as the peak of the intron counts did not precede the exon counts, indicating a lack of splicing or an mRNA that was as unstable as the intron. Using the Ensembl browser we determined that this could be due to previously detected alternatively initiated or spliced mRNAs and these two profiles were also removed from the analysis (Per1 and Myl12a). This left 25 serum response gene expression profiles for analysis.

### RNA ligation-mediated poly(A) test

The RL-PAT was an improved version of a previously reported method [52], in which an oligo is ligated to the end of the mRNA and its complementary oligo is used to prime reverse transcription. A gene specific primer is then used in combination with the reverse transcription oligo to amplify a section of 3’ UTR with the attached poly(A) tail. The efficiency of the ligation reaction was improved by using a mutated ligase that needs a 5’ to 5’ adenylated substrate. The mutated ligase is incapable of performing the 5’ to 5’ adenylation that is the intermediate step in the catalysis of nucleic acid ligation, and therefore cannot ligate cellular RNAs to each other, but only to the pre-adenylated anchor oligo. The RL-PAT is graphically explained in Supplementary Figure 10. Details of the RL-PAT are as follows: The 5’ to 5’ adenylated and 3’ blocked ‘PAT anchor’ oligo was ligated to the 3’ end of total RNA overnight at 16°C using RNA ligase 2, truncated KQ (NEB, M0373). The ligated RNA was reverse transcribed using SuperScript III Reverse Transcriptase (Invitrogen, 18080044) with the ‘PAT-R1’ oligo (complementary to ‘PAT anchor’) to form cDNA. This cDNA was then used for PCR with GoTaq G2 Flexi Polymerase (Promega, M7801), using a forward primer annealing to the 3’ UTR of the mRNA of interest (all primer sequences in Supplementary Table 5) and PAT-R1 as the reverse primer. All mRNA specific PAT primers were validated by performing PAT on mRNA deadenylated with oligo-d(T) and RNAse H (Supplementary Fig. 11) and PAT products were sequenced to ascertain their identity. In addition, we compared two mRNA specific PAT primers for the 5 constitutively expressed mRNAs and found the poly(A) tail distributions were virtually identical (Supplementary Fig. 12). A detailed protocol is available upon request. A more sensitive, nested version of this method has now also been published [53].

### Gel electrophoresis, scanning and sizing of PAT products

PCR products were run alongside 100bp ladder (NEB, N3231L) on a 1.2% agarose gel prestained with SYBR Safe (Invitrogen, S33102) and recorded using a ChemiDoc (Bio-Rad) or a LAS-4000 (Fujifilm), taking care not to overexpose any bands. Intensity profiles were generated by quantitative gel scanning analysis. Lanes were scanned in the Quantity One software package (Bio-Rad, 1709602), background lane intensity was subtracted, the intensity corrected for the size of fragments, and the data normalised to the area under the profile. The 100bp ladder was used to map distance migrated to length in base pairs using a fitted polynomial curve. The predicted fully deadenylated product size was subtracted from the profiles to obtain profiles corresponding to the poly(A) tail sizes alone. Supplementary Fig. 13 shows this method applied to gels with restriction digests and demonstrates that individual band sizes determined this way are within 10 nucleotides of the actual size in 95% of the cases within the 250-500 nucleotide range that we require for PAT. Averages of three or more replicates are within 3 nucleotides of the actual size in 95% of cases, demonstrating that the gels effectively measure fragment size in the desired range. Errors are larger for fragments under 200 base pairs or over 550 basepairs.

### Thiouridine labelling

Cells were cultured as indicated and 4-thiouridine was added to a concentration of 250 μM from a 250 mM stock in DMSO for the indicated times. RNA was isolated using TRI-Reagent (Sigma). Biotinylation and isolation of labelled RNA was essentially as described by Rabani et al [1]. Further detail on thiouridine labelling during the serum response can be found in Supplementary Figures 7 and 8.

### LC-MS/MS analysis of intracellular 4-thiouridine and 4-thiouridine triphosphate

Following PBS washing, cells were directly extracted in situ with ice cold methanol containing mM EDTA. Analysis was carried out on an Agilent 1100 HPLC system (Agilent, Cheshire, UK) coupled to a Waters Quattro Ultima (Elstree, UK) triple quadrupole mass spectrometer in negative electrospray ionisation mode. Separations were achieved on a Luna C-18, 3 μm (2.1 × 150 mm) column with Security Guard (4 × 2 mm) at 40°C (Phenomenex, Cheshire, UK). The standards and the samples (5 μl injection volume) were eluted at a flow rate of 200 μL/min using a gradient mobile phase containing of 5 mM dimethylhexylamine (DMHA) in water: methanol (95:5 v/v) (A), and 5 mM DMHA in methanol: water (80:20 v/v) (B). The MS source temperature was at 125°C with nitrogen as drying and nebulising gas with precursor ions of *m/z* 258.92 and *m/z* 498.90 and product ions of *m/z* 126.20 and *m/z* 401.10 respectively for 4-thiouridine and 4-thio-UTP. Concentrations were determined by the internal standard method with extracted calibration standards using an average measured intracellular volume of 4.7 pL and data analysis was done using Waters MassLynx™ Software.

### siRNA knockdown in NIH-3T3 cells

Cells of passage 20 or lower were seeded in DMEM containing 10% new born calf serum at a density of 9.2×10^3^ cells/cm^2^. 24 hours later, media was refreshed and siRNA (SMARTpool ON-TARGET plus, Dharmacon) was added to 10nM using Lipofectamine RNAiMax and Optimem according to the manufacturer’s instructions. This was repeated after a further 24 hours. On day 4, cells were washed with PBS and media replaced with 0.5% serum DMEM. 24 hours later cells were serum stimulated by adding serum to 10%.

### Western blotting

Protein samples were prepared for whole cell lysate using RIPA buffer and SDS loading buffer was added. For cytoplasmic and nuclear fractions, SDS loading buffer was added to the supernatant and pellet fractions prepared as described under RNA isolation. Loading volumes were chosen such that either cell number (fractionation) or total protein content (Cnot1 knockdown) was consistent. SDS-page gel electrophoresis was performed using either 8% (Cnot1) or 12% (fractionation validation) acrylamide and a constant current of 20mA per gel. Proteins were transferred to PVDF using the semi-dry transfer method and detected by western blotting using the following antibodies: Cnot1 Polyclonal antibody (Proteintech), Lamin A/C clone 4C11 (Cell Signalling), α-tubulin 236-10501 (Molecular Probes).

### Mathematical models and fitting

#### Introduction to mRNA decay models

The basic building block of the models can be described in the following manner:

1. An unspliced non-parametric function *p*(*t*) is derived from the unspliced data levels using a modified Gaussian process.
2. The unspliced mRNA level *p*(*t*) is used as the input for an ordinary differential equation *(ODE)*-level model for the mature mRNA levels.
3. The parameters of the model (i.e. α, λ, τ) are fitted by minimising the difference between the model prediction and the data.

#### mRNA decay models

The two models used for the mature mRNA data are as follows:

##### Immediate decay model

The immediate decay model (ID) follows classical exponential decay. As in all models the changes to the mRNA level (*m*;) are dependent on the splicing of the precursor (*p*) with a splicing rate *α*. In this model the newly made mRNAs are then immediately available for decay with a decay rate *λ*. Each mRNA has the same chance *λ* of decaying regardless of its age. The average lifetime of an mRNA in this model is 1/*λ*:

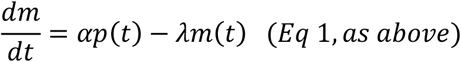

##### Delayed decay model

In the **delayed decay model (DD)** all mRNAs are protected from decay for a fixed time *τ* after their splicing, becoming susceptible to decay after this period has passed. This model assumes that, although there may be a large number of catalytic steps that postpone decay, all with similar rates *k*_1_ to *k*_*n*_, these can be modelled as a single delay. Each catalytic step will take, on average 1/*k*_*i*_, and so by the Central Limit Theorem, the distribution of total delay time will converge in mean to the sum of the mean times for each step, assuming similar behaviour for individual mRNAs transcribed from the same sequence:

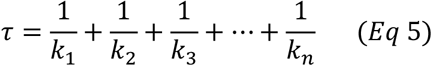

The deadenylation of a poly(A) tail is a clear example where such an approximation is likely to apply to due to the collective effect of the removal of 200-250 individual adenosines. If their rates are slow enough to contribute to the delay in decay, but not much slower than the deadenylation rate, this delay could also include effects of export, uridylation and decapping. In this model there are two pools of mRNA, a protected pool *m*^*P*^, which is constituted of the mRNAs synthesised during the preceding time *τ* and a susceptible pool *m*^*S*^ in which the mRNAs that are older than *τ* are subject to exponential decay with a rate *λ*. For this model the chance of decay for each mRNA changes from 0 to *λ* at age *τ*. The average lifetime of an mRNA in the DD model is *τ* + 1/*λ*:

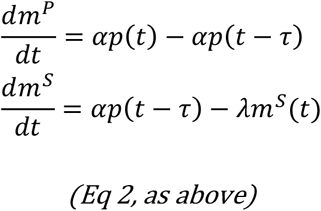

#### Fitting of the decay models

For the qPCR data, we have measured unspliced and spliced mRNA levels directly. In this case, as any measurement error is likely to be in the difference in C_t_ values, for which the natural measure of errors is in log space, our fitting residual sum of squares between the logarithm of the data mature mRNA levels and the logarithm of the model mature mRNA levels (for models with two pools of mature mRNA, the model mRNA is taken to be the sum of the two pools):

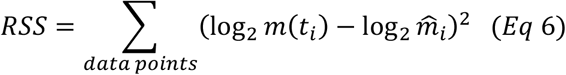

For the NCounter data, the data are given in terms of intron and exon levels, measured by specific probes from Nanostring. In this case, the unspliced levels for the model are given by the relevant intron level (i.e. the input *p* knots for the modified Gaussian process are given by 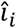). Fitting is then to the exon levels, which in this case map directly to the sum of the unspliced and mature mRNA levels in the model. In order to model the potentially different intron and exon probe efficiencies in the NCounter procedure, the residual sum of squares is given by:

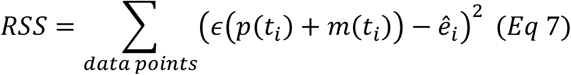

where *ϵ* is the ratio between the exon and intron probe efficiencies. Note that this extra parameter *ϵ* is a feature of the measurement method, and so is independent of the serum time course and independent of the model fitted. In practice this means that the four combinations of the two time courses and two models for each mRNA are fitted simultaneously, ensuring that the fitted *ϵ* is the same in all cases.

#### Modifying the decay model for changing delays

Starting from the equation for the protected class in the delayed decay model:

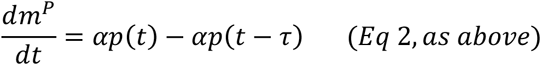

the level of the protected class of mRNA at time *t* can be written

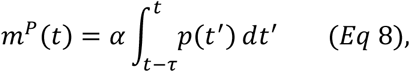

plus a constant dependent on the initial conditions. An interpretation of this integral is that the level is equal to all pre-mRNA spliced between times *t* − *τ* and *t*. If the delay changes over time, this integral can be re-written in terms of the time *g*(*t*) at which mRNA losing protection at time *t* (whether through completion of deadenylation, uridylation or other marking for decay) was originally produced through splicing:

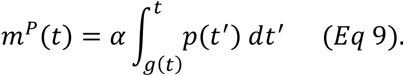

This leads to the following system of differential equations for more general changing delays:

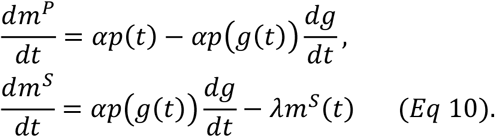

Special cases of this system are the immediate decay model *g*(*t*) = *t* and the delayed decay model *g*(*t*) = *t* − *τ*, which give the same differential equations as mentioned in the main text (note that for the immediate decay model, 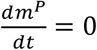, so there is no protected class).

In general, the level of susceptible mRNA *m*^*s*^(*t*) can be calculated by solving the differential equation, given the function *g*(*t*), and then *m*^*p*^(*t*) can be found from the integral.

Figure 6 was generated assuming the following form for *g*(*t*):

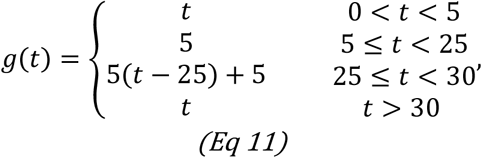

and assuming a Gaussian pulse in the unspliced level, corresponding to a delay increasing from 0 to 20 at time *t* = 5 minutes, then dropping back to 0 by time *t* = 30 minutes. This is compared against a simulation of the delayed decay model with a constant delay of 20 minutes.

#### Poly(A) tail model

The poly(A) tail model used here can be constructed from three main assumptions:

1. The poly(A) tail distribution of newly-spliced RNA is independent of the time when it is produced. The level of newly-spliced RNA is taken to be proportional to the intron level as derived from the NCounter data.
2. Poly(A) tails are deadenylated at a tail-dependent rate which can be decomposed as a maximum rate with a length-dependent factor between 0 and 1. This models the deadenylation enzyme losing its binding as the tails get shorter.
3. Once poly(A) tails are short enough, the body of the mRNA is degraded by an exonuclease at a tail-dependent rate, also writable as a maximum rate constant times a length-dependent factor between 0 and 1, showing increased binding of the decay enzyme as tails get shorter. Once an mRNA is bound by an exonuclease, it is assumed to be instantly removed from the distribution.

The factors affecting the amount of mRNA with a tail of given length *i* are deadenylation from longer tails, deadenylation to shorter tails, decay and input from newly-spliced RNA.

If *y*_*i*_(*t*) is the amount of mRNA with a given tail length *i*, an ordinary differential equation can be written to express the rate of change of this concentration, given the deadenylation 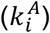 and decay 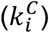 rates at that length, and the rate of mRNAs of this tail length being produced (*Q*_*i*_(*t*)) as

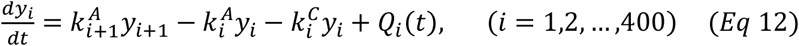

The first two terms of this differential equation arise from deadenylation, firstly from tails of length *i* + 1 down to length *i*, and then from tails of length *i* to *i* − 1, under the assumption that the enzymes deadenylate one nucleotide at a time. The third term of this equation represents mRNA decay, and the final term represents the input of mRNA at that length. This differential equation holds for all lengths *i* considered in this model, from 1 adenosine residue up to a possible maximum length of 400.

The dynamics of the model is completely described using two reaction constants (*k*^*A*^, *k*^*C*^), and four normal distribution parameters (*μ*_*C*_, *μ*_*A*_, *τ*_*C*_, *τ*_*A*_). The constants 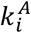 and 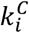 for each length are given by

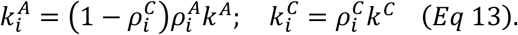

The rate constants are: *k*^*A*^ for deadenylation, and the decay rate *k*^*C*^. Variations in deadenylation and decay rates at each length are given by the rho functions 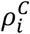, representing the fraction of mRNAs of length *i* bound to the decay enzyme, and 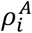, representing the fraction bound to the deadenylation enzyme. A further assumption, relating the mean and standard deviation of length at which the degrading enzyme is bound (*μ*_*C*_ and *σ*_*C*_), and the mean and standard deviation of the length at which the deadenylator unbinds (*μ*_*A*_ and *σ*_*A*_), means that these rho functions are written in terms of the cumulative normal distribution function *ϕ*.

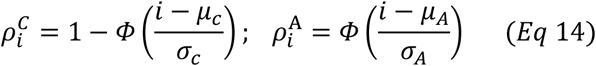

Mathematical analysis of this set of differential equations (one for each length, Equation 14), demonstrates that if no input or decay is occurring, then the speed at which a band of mRNA moves along the gel (in bases per minute) is equal to the deadenylation rate for those lengths. Using this fact, we calculated a model-estimated average deadenylation time using the best-fit deadenylation rates via the following formula

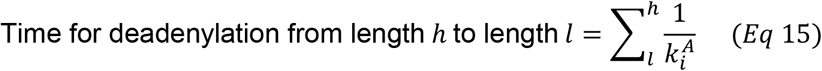

#### Input functions of the poly(A) model

The input function *Q*_*i*_(*t*) is given by a shape *q*_*i*_ independent of time multiplied by a time envelope *f*(*t*). The time envelope is taken to be proportional to the level of unspliced mRNA, the non-parametric interpolation function *p*(*t*), derived from the NCounter intron data.

The input tail distribution for this work was assumed to be Gaussian in shape – there is insufficient data on the control of poly(A) tail length in the nucleus to suggest a more precise shape, we therefore choose to make the simplest assumption regarding the poly(A) tail distribution as

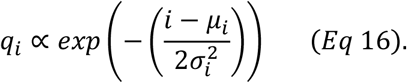

##### Fitting details: poly(A) tail model

As the PAT assay is not quantitative (it only returns normalised distributional data), a different approach was required for fitting the model – we found that a naïve ordinary least squares approach led to the optimisation not converging to a sensible tail distribution prediction.

The function minimised can be written succinctly as

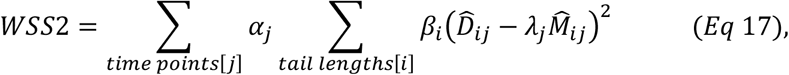

with time weights *α*_*j*_, length (quality) weights *β*_*i*_, and scaling factors *λ*_*j*_.

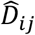 and 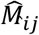 are transformed data (at time point *j* and length *i*) and model predictions respectively. The forms of these parameters is discussed below. The procedure for calculating this expression is broken down into the main steps:

1. We transformed the data and the model. Firstly, at each time point *j*, we scaled the data and the model so that their maximum values are 1, via

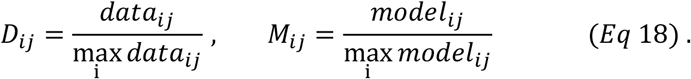 Secondly, we stretched the data to give more weight to the highest intensity values (i.e. those close to 1), where the PAT PCR product well above any background noise, via

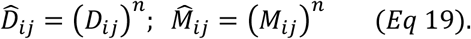
2. We calculated the time scaling factors, *λ*_*j*_, by minimising the least squares difference between the transformed data and the transformed model (at time *j*), weighted by the length quality weights *β*_*i*_ (the inner sum above), namely

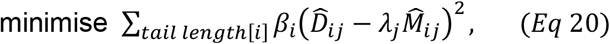

giving

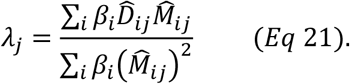
3. Finally, we summed the inner expression of (Equation 21) over all time using the time weights *α*_*j*_ (given by the mature mRNA levels, estimated directly from the difference between the exon and intron levels in the NCounter data, not using any information from our decay modelling).

The weights *α*_*j*_, *β*_*i*_ were chosen to give more emphasis in the sum to the position of the high intensity band through the stretching transformation and to the data available during the period of highest mRNA concentrations as the PAT assay is more accurate when more mature mRNA is present in the analysed sample, due to the PCR required as part of the method. *α*_*j*_ is the mature mRNA level at time point *j* and *β*_*i*_ was set to 1 for lengths below 300 nucleotides, and 0 otherwise.

### Protocol, data and script availability

Detailed protocols for RL2-PAT, thiouridine labelling and nuclear and cytoplasmic RNA isolation as well as the NCounter data, the quantification of the PAT gels and the scripts used in analysing and modelling these data are available upon request.

## SUPPLEMENTAL MATERIAL

Supplemental Figures and Tables are available for this article

## ACKNOWLEDGMENTS

Peter Shaw, Keith Spriggs, Tillman Achsel and Sebastiaan Winkler are thanked for critical reading of early versions of this manuscript. Irham Fatema Goulamaly and Che Ku Norhafizan Che Ku Salim are thanked for some initial observations on poly(A) tail changes during their MPharm research projects in the spring of 2012.

## FUNDING

This work was supported by the Biotechnology and Biological Sciences Research Council [grant numbers BB/G001847/1, BB/K008021/1, BB/J014508/1]. Hannah Parker, Raj Gandhi and Kathryn Williams received BBSRC funded PhD studentships. Clara Daher was an Erasmus exchange student from the University of Rennes 1 during the spring of 2015.

## AUTHOR CONTRIBUTIONS

CdM had the original idea and supervised the experimental work. HNP and ABB generated important pilot data. RS performed the majority of the experiments. RDG, CD, KW and CdM contributed to the 5-minute serum time course data. RDG developed the RL2-PAT. KW checked PAT products by deadenylation and sequencing and performed the Cnot1 knockdown and chromatin associated RNA experiments. WU determined thiouridine and thioUTP levels in cells, supervised by DAB. MA-S checked gel resolution and tested different PAT primers for the same mRNA as shown in the supplement. PAT test gels were digitised, calibrated and processed by RDG, KW and RS. JADW and GJT developed models which were implemented in R (R Consortium, 2019) by GJT. JADW and GJT discussed the best ways of optimising the fitting procedure. GJT, RS and CdM wrote the manuscript.

