## Supplementary Figures for "Nuclear poly(A) tail size is regulated by Cnot1 during the serum response"

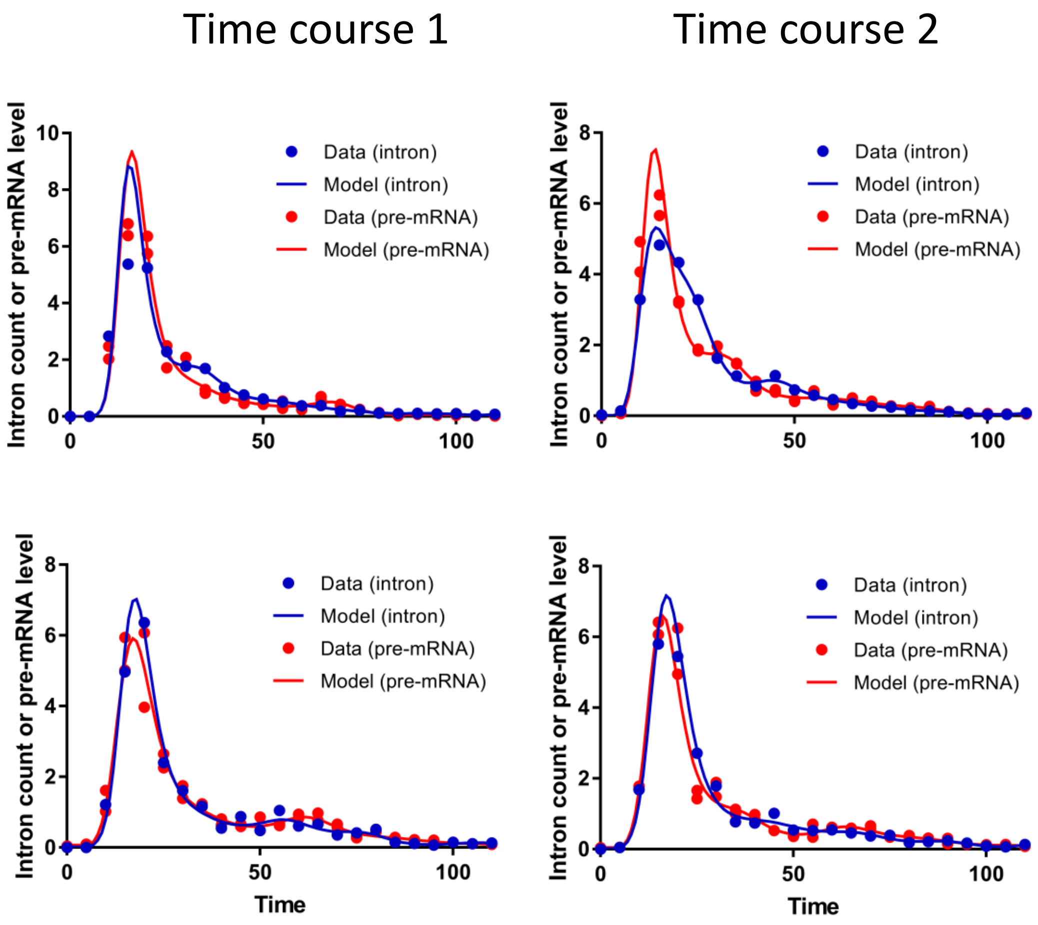


Supplementary Figure 1. **Comparison of Fos and Fosb pre-mRNA levels determined by RT-qPCR and Ncounter**. Data (dots) are shown with the non-parametric fitted profiles used in the models (lines).

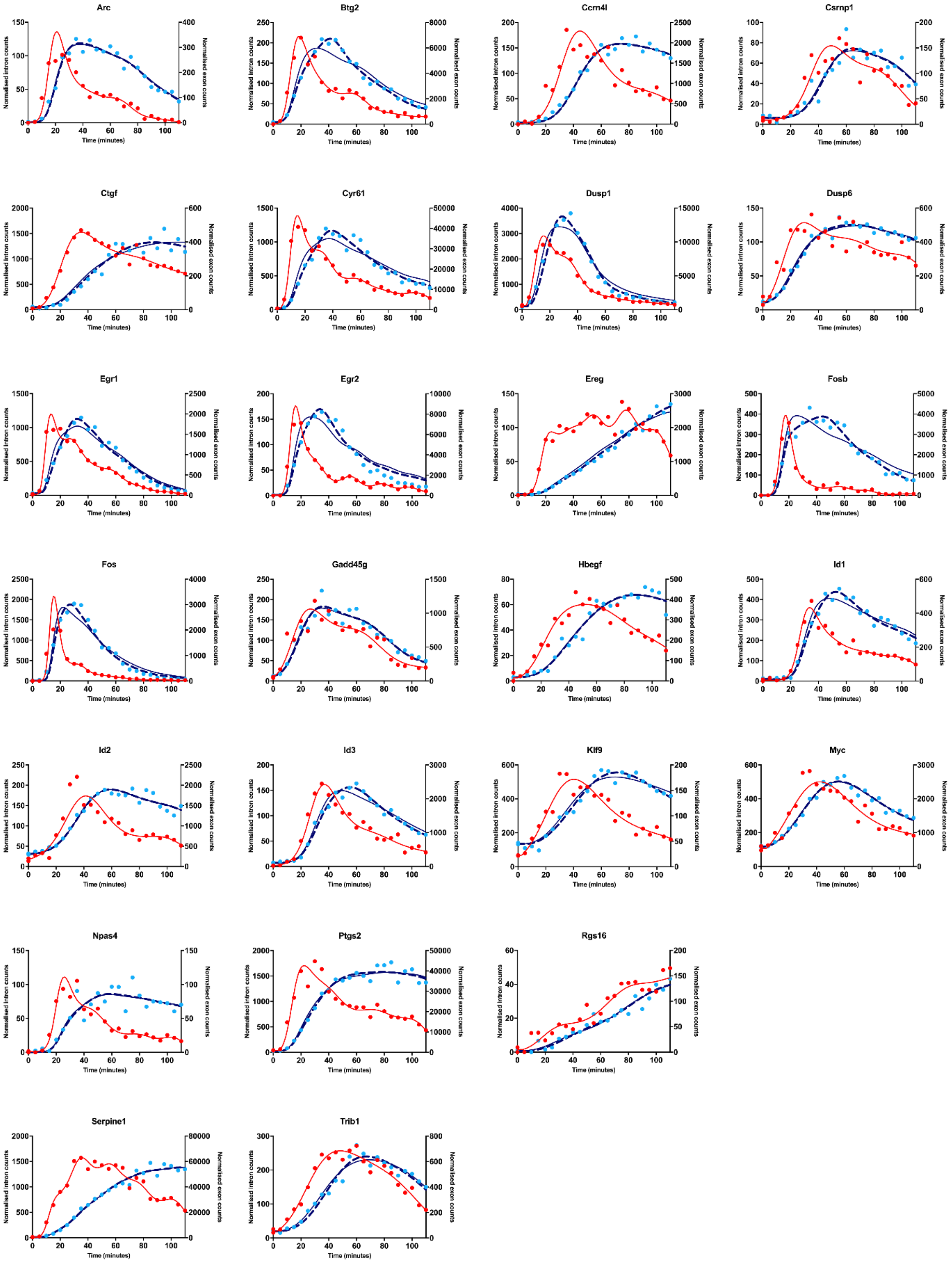


Supplementary Figure 2A. **NCounter data for serum time course 1 fitted with immediated and delayed decay models.** As in Figure 3, intron counts (pre-mRNA) are in red, exon counts (mRNA + pre-mRNA) are in blue. Dots are data, lines are models fitted to the data. The continues blue line show immediated decay and the interrupted blue line shows delayed decay models.

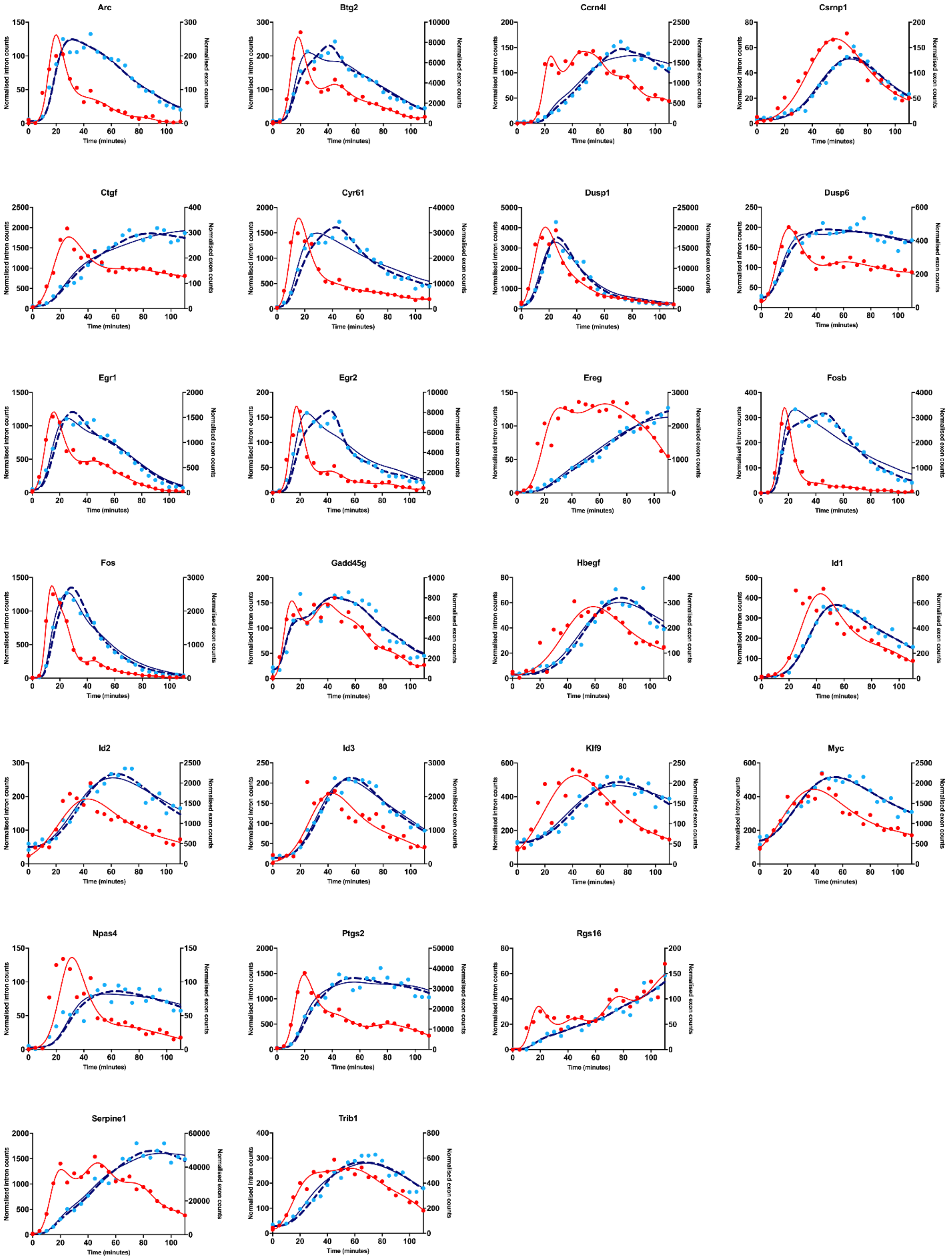


Supplementary Figure 2B. **NCounter data for serum time course 2 fitted with immediated and delayed decay models. .** As in Figure 3, intron counts (pre-mRNA) are in red, exon counts (mRNA + pre-mRNA) are in blue. Dots are data, lines are models fitted to the data. The continues blue line show immediated decay and the interrupted blue line shows delayed decay models.

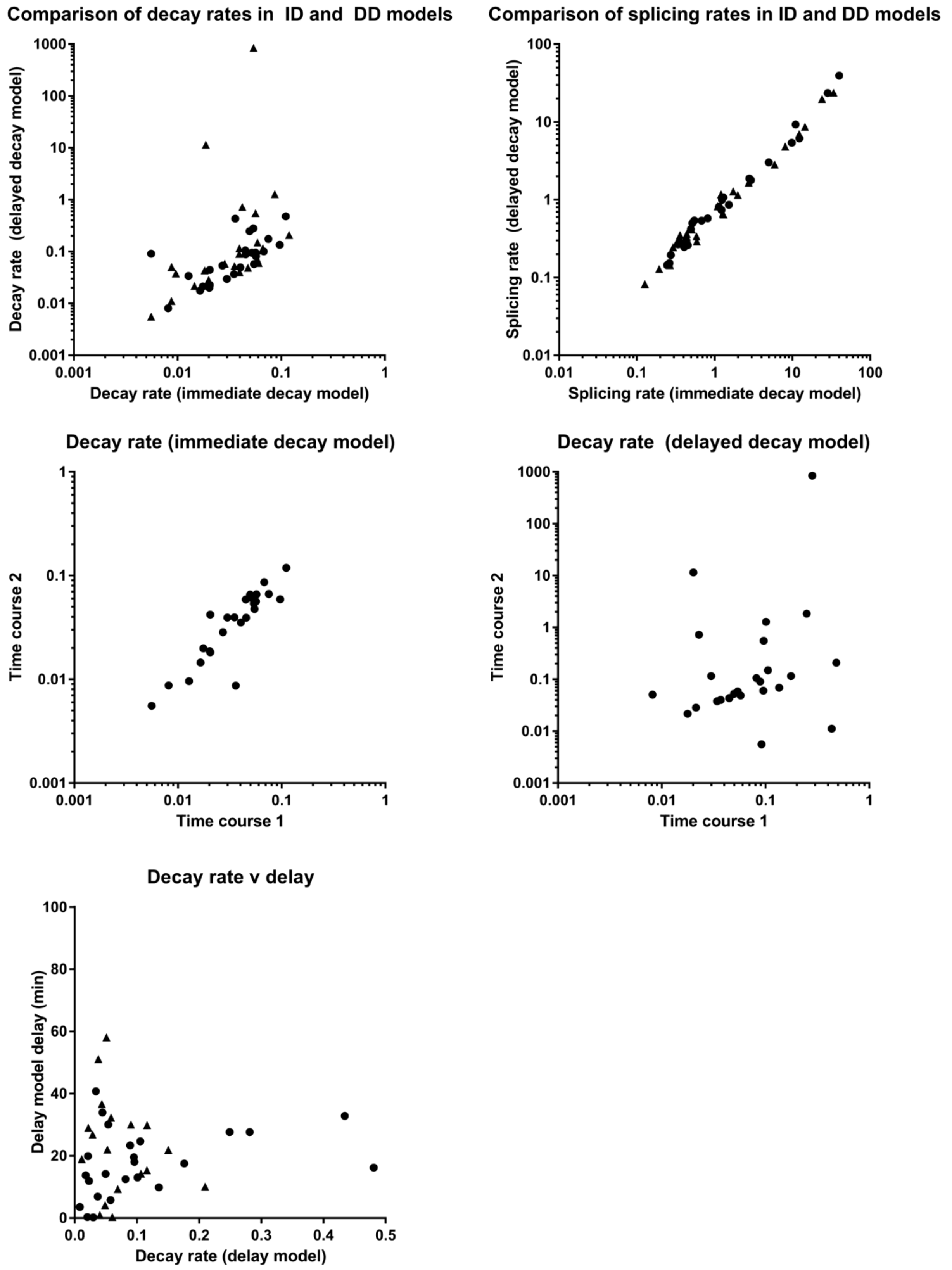


Supplementary Figure 3. **Comparisons of the fitted parameters for the modelling of the NCounter data.** Where both are shown, circles indicate Time course 1 data and triangles indicated Time course 2 data.

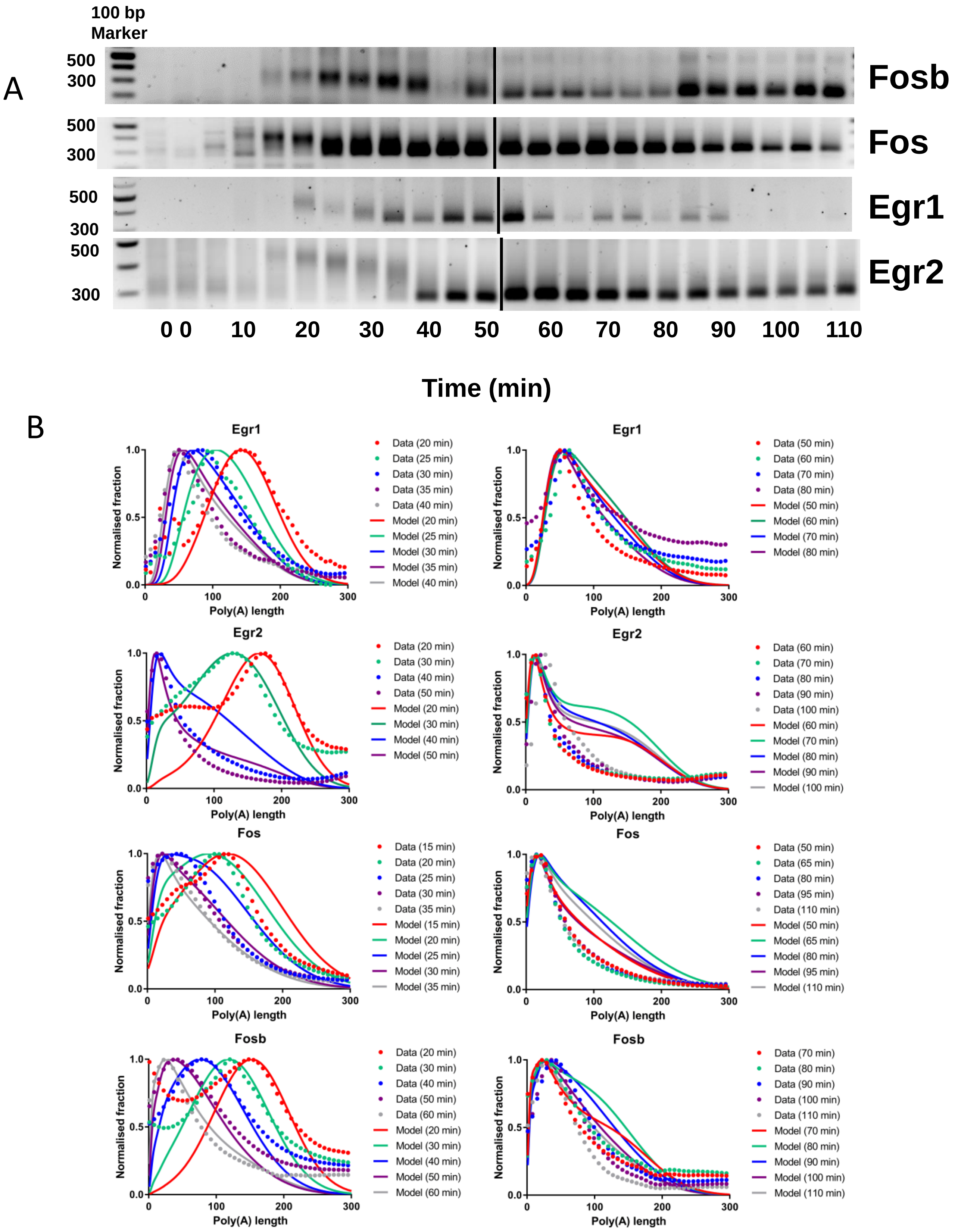


Supplementary Figure 4. **Additional poly(A) modelling data for serum time course 1**. A. For details see Fig 5A B. For details see Fig 5

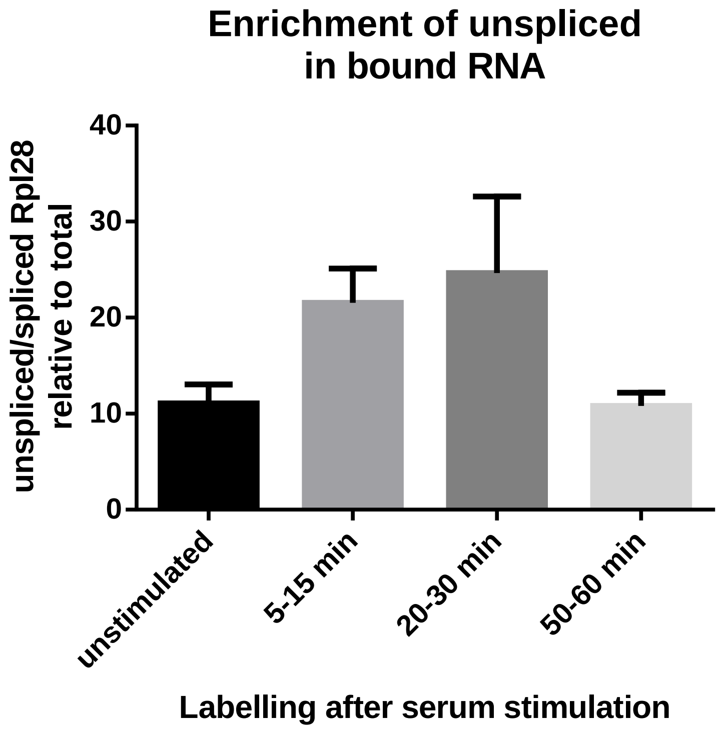


Supplementary Figure 5. **Enrichment of unspliced mRNA in the bound fraction of thiouridine labelled RNA**. An aliquot of the RNA used for PAT in Figure 6A was reverse transcribed using random primers and used for quantitative PCR with primers for spliced and unspliced Rpl28 mRNA.

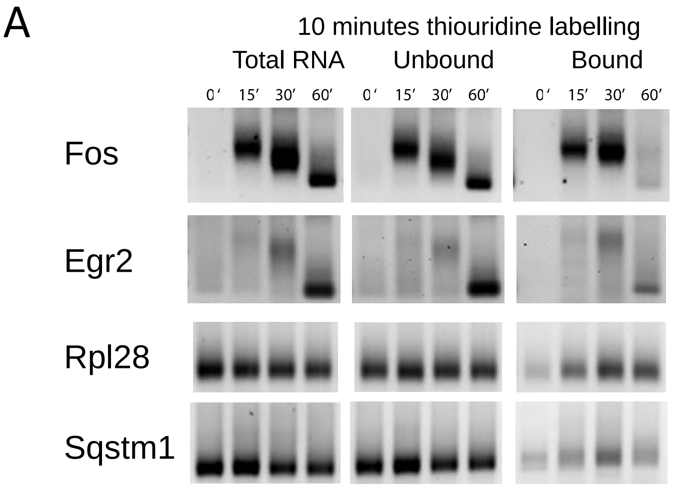


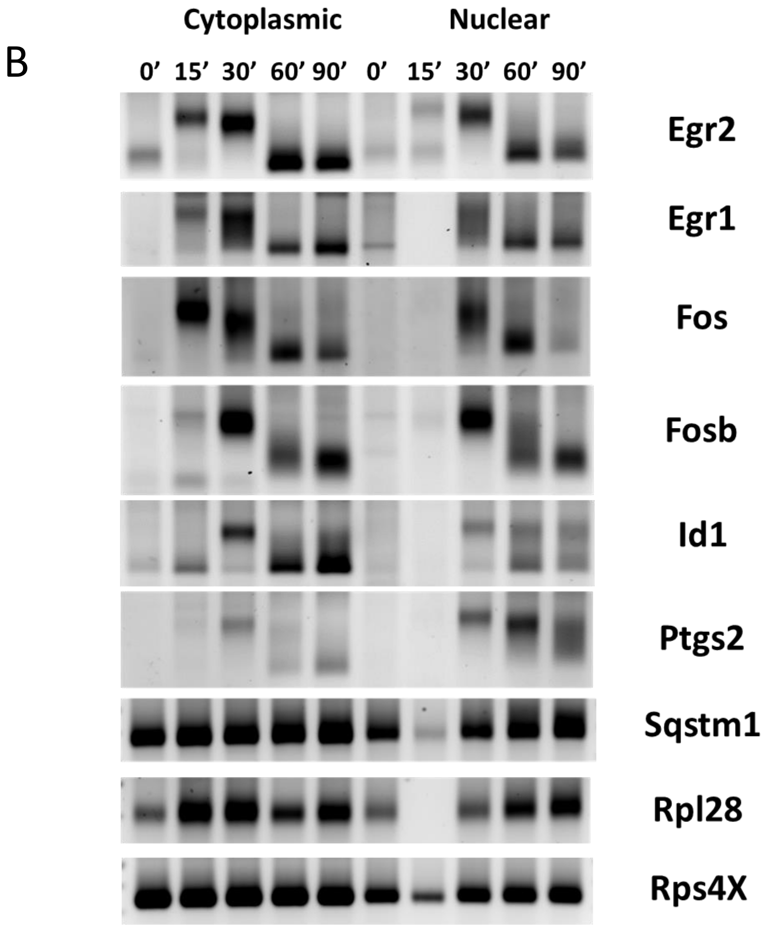


Supplementary Figure 6. **Initial poly(A) tail size is regulated**. A.NIH3T3 cells were serum starved and incubated with 4-thiouridine for 10 minutes, either without stimulation with serum (0') or with serum addition from 5 to 15 minutes (15'), 20 to 30 minutes (30') or 50 to 60 minutes (60'). Total RNA was isolated and labelled RNA was isolated from a fraction (bound). RL2-PAT was performed for the indicated mRNAs. These are independent replicates of Figure 5.

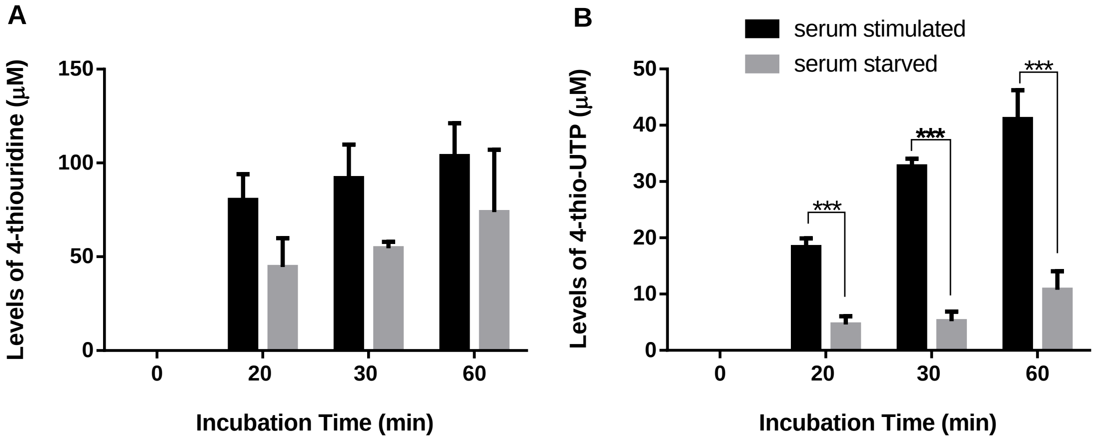


Supplementary Figure 7. **Accumulation of 4-thiouridine (A) and 4-thio-UTP (B) in NIH 3T3 cells** that were incubated with 250 μM 4-thiouridine for the indicated times. Each bar represents the mean ± SEM from 3 separate experiments. ****= $p < 0.0001$; ***=$p < 0.001$; **= $p < 0.01$; *= $p < 0.05$. These data explain the consistently low levels of bound mRNA we obtained in unstimulated cells, which made it impossible to determine initial poly(A) tail sizes of serum response mRNAs before induction in Figure 5A. Serum stimulation is probably increasing the phosphorylation of 4-thiouridine.

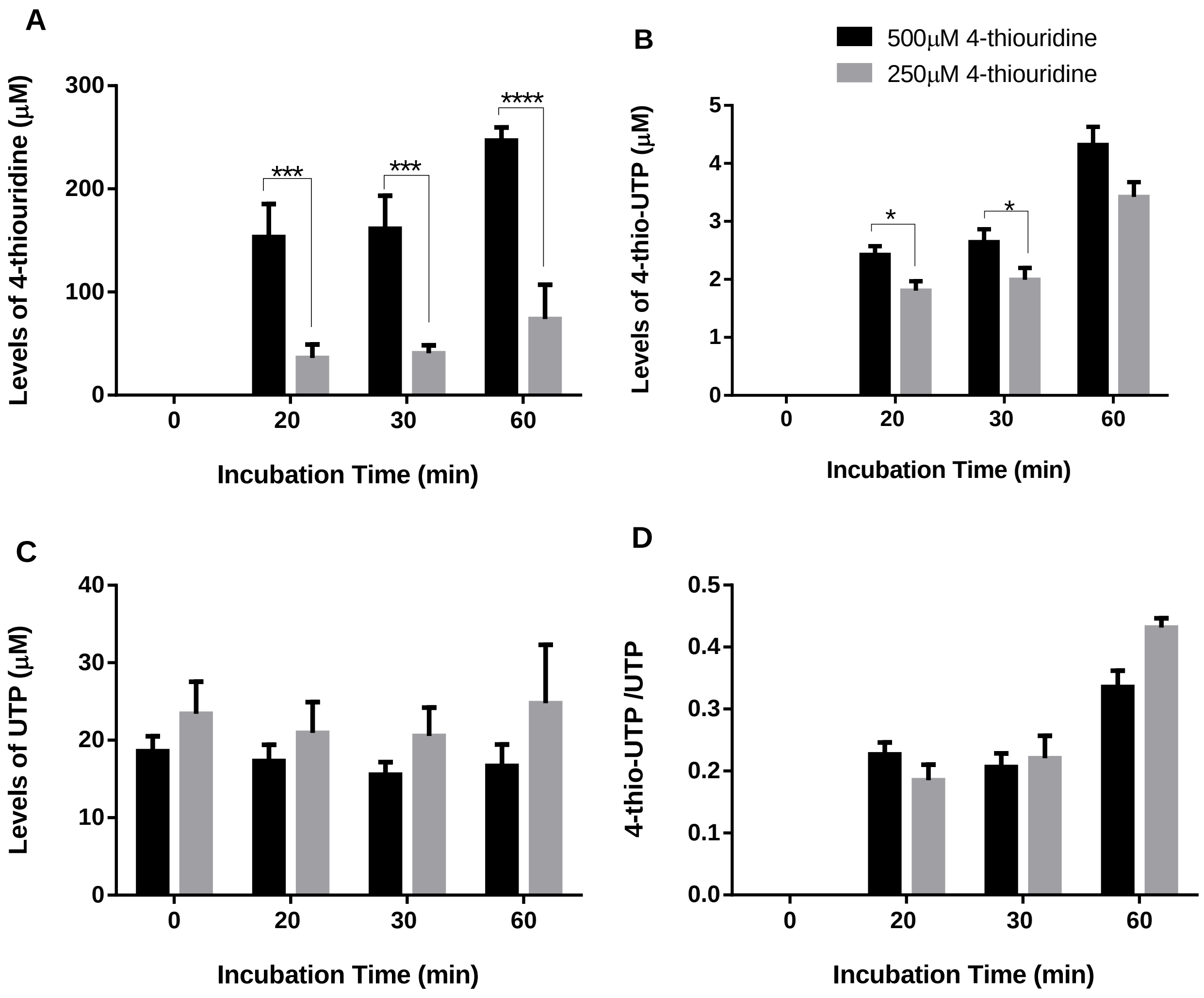


Supplementary Figure 8. **Higher thiouridine concentrations do not increase intracellular thioUTP levels.** Levels of thiouridine (A), 4-thio-UTP (B), UTP and the ratio of 4-thioUTP/UTP (D) in NIH 3T3 cells that were incubated with either 500 µM or 250 µM 4-thiouridine for the indicated times in cells seeded the day before (not serum starved or stimulated). Each bar represents the mean ± SEM from 3 separate experiments. ****= $p < 0.0001$; ***=$p < 0.001$; **= $p < 0.01$; *= $p < 0.05$.

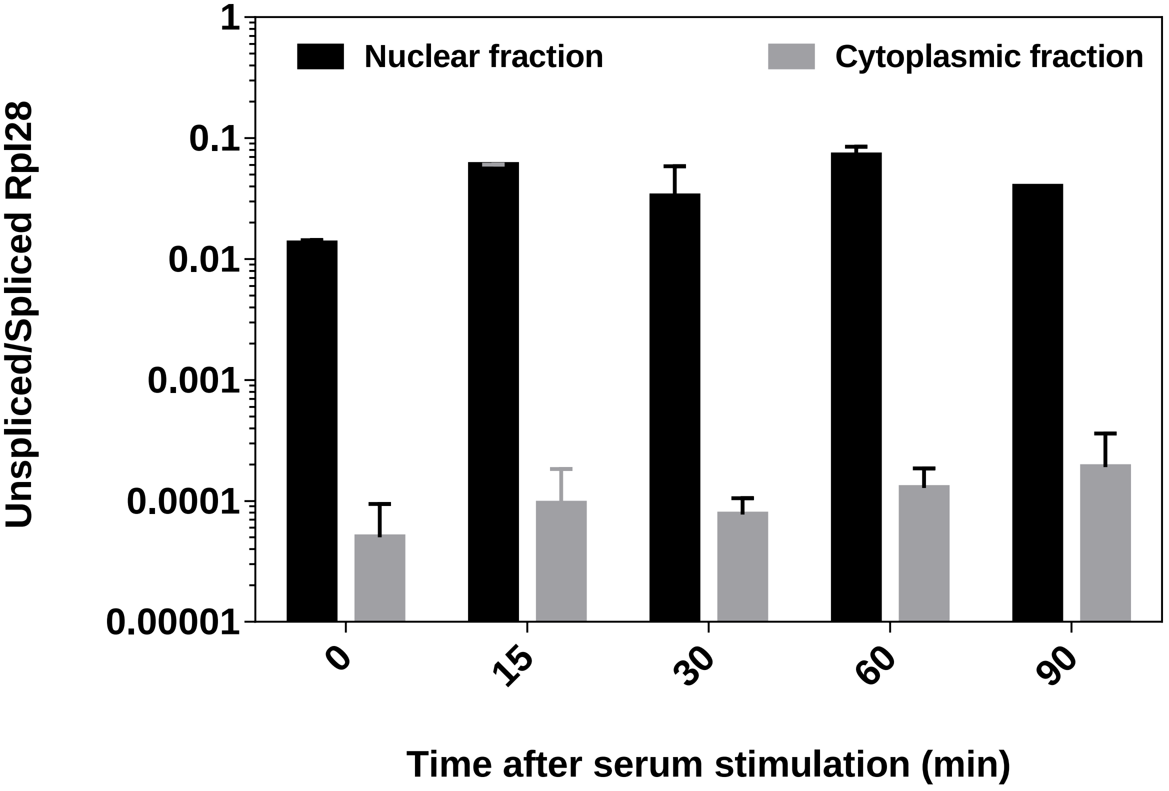


Supplementary Figure 9. **Enrichment of unspliced mRNA in the nuclear fraction.** An aliquot of the RNA used for PAT in Figure 6B was reverse transcribed using random primers and used for quantitative PCR with primers for spliced and unspliced Rpl28 mRNA.

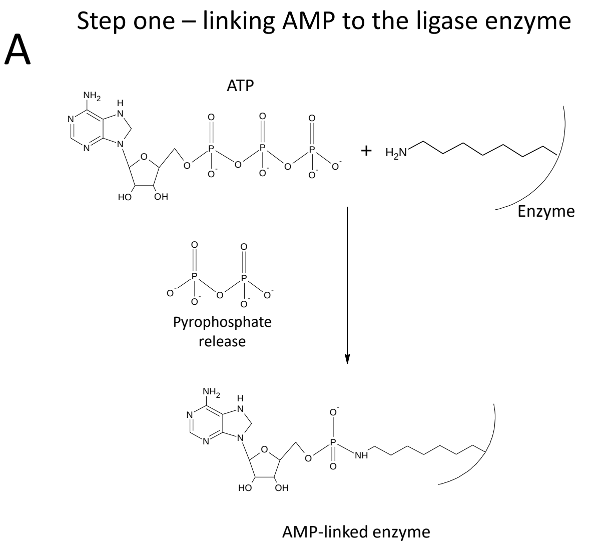


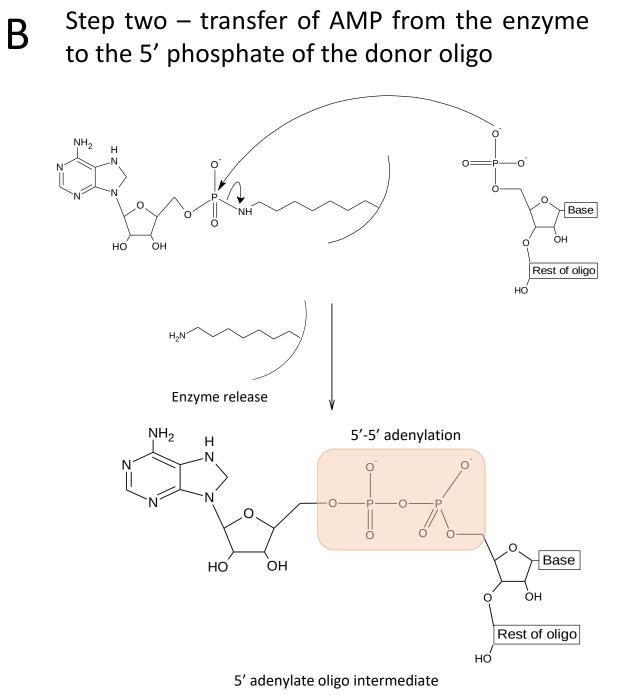


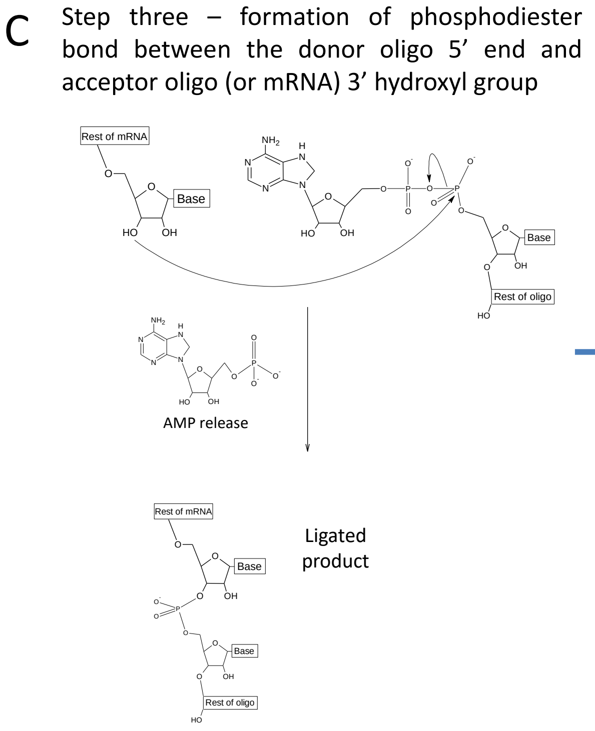


| Enzyme | Step 1 | Step 2 | Step 3 |
|---|---|---|---|
| RNA ligase 1 | ✓ | ✓ | ✓ |
| Mth RNA ligase | ✓ | ✓ | × |
| Rnl2 truncated | × | × | ✓ |

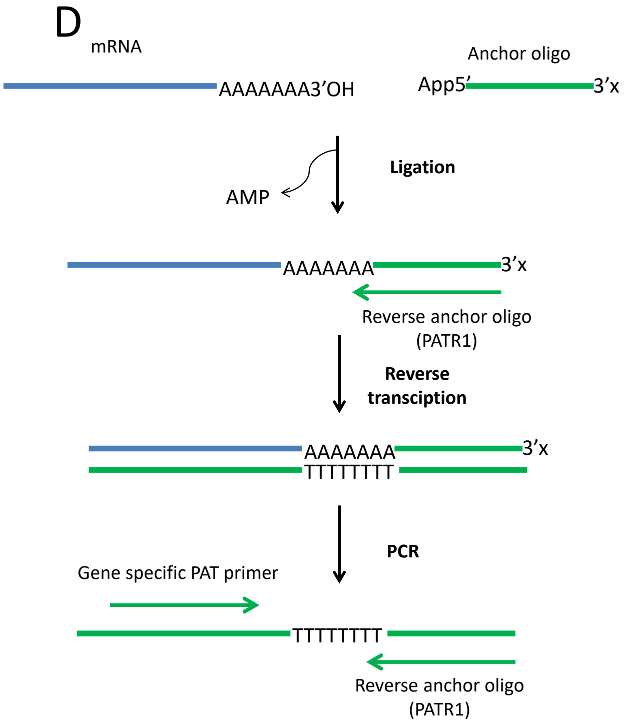


Supplementary Figure 10. **Diagrams explaining the RL2 PAT**. (A-C) intermediary steps affected in available RNA ligase 2 mutants (D) the principle of the RL2-PAT.

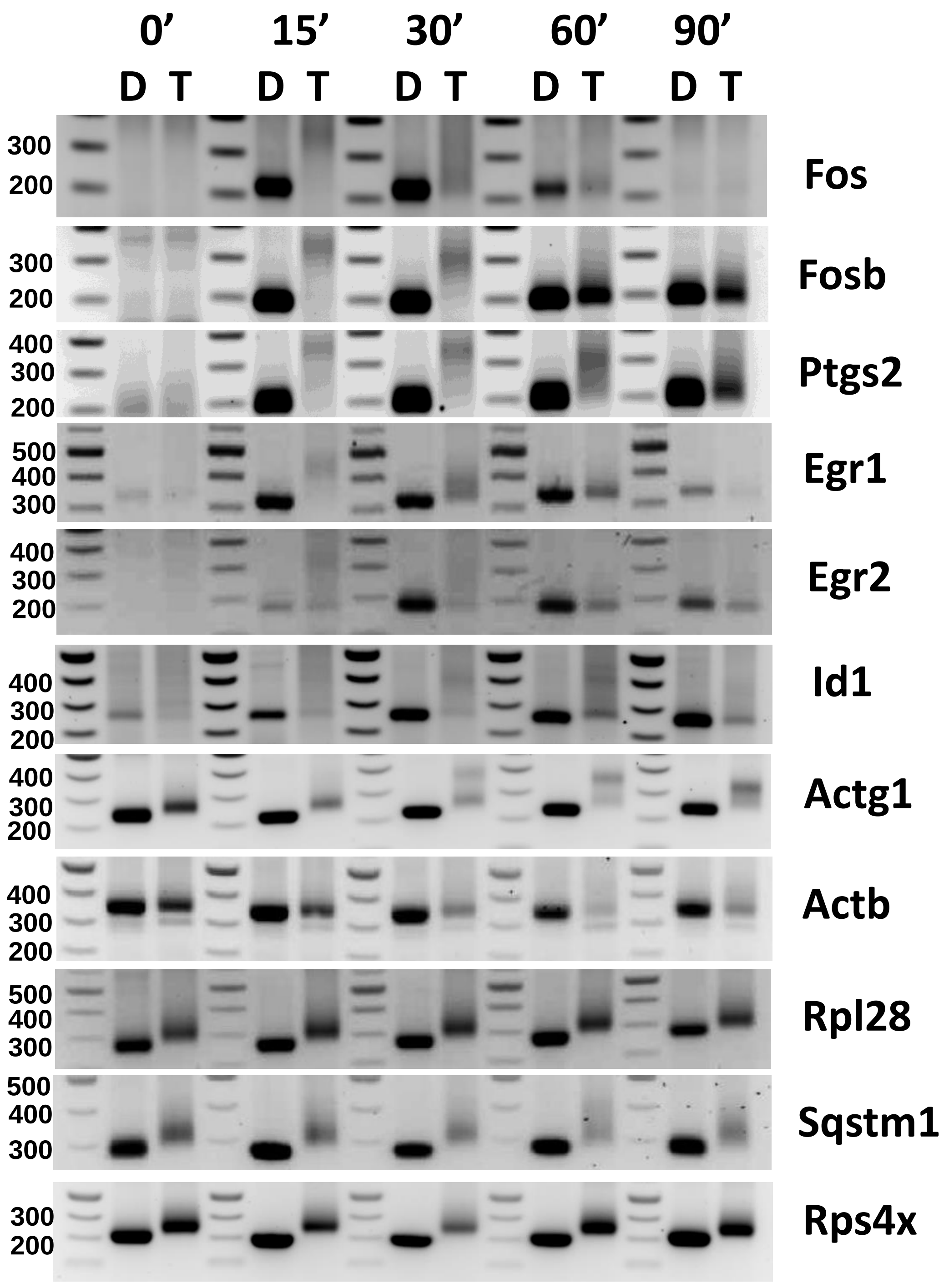


Supplementary Figure 11. **Changes in poly(A) tail size are not due to alternative polyadenylation**. Serum starved NIH3T3 cells were treated with serum for the indicated time periods and RNA was isolated. LR2-PAT was performed for the indicated mRNAs on total RNA (T) and RNA treated with RNase H and oligo dT (D). Sizes of the 100 bp marker are indicated.

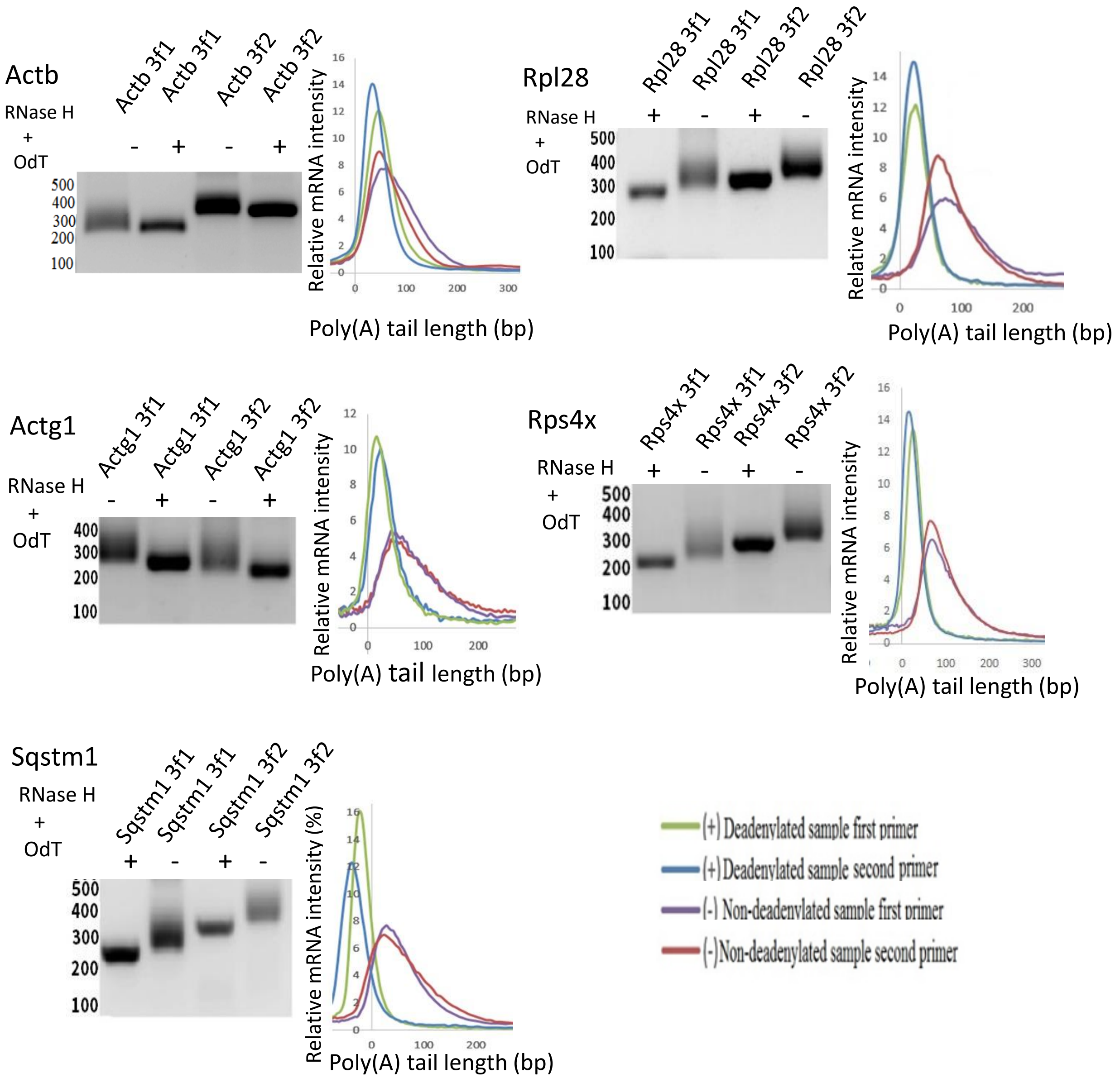


Supplementary Figure 12. **RL2-PAT PCR with different primers for constitutively expressed mRNAs gives the same poly(A) tail distributions.** Total RNA was isolated form NIH3T3 cells seeded 24 hours previously (no serum starvation). RNA was treated with RNAseH and oligodT (+) or not (-). RL2-PAT was performed for the indicated mRNAs using 2 different PAT primers and processed using our quantitative gel scanning method. Normalised intensity profiles are shown.

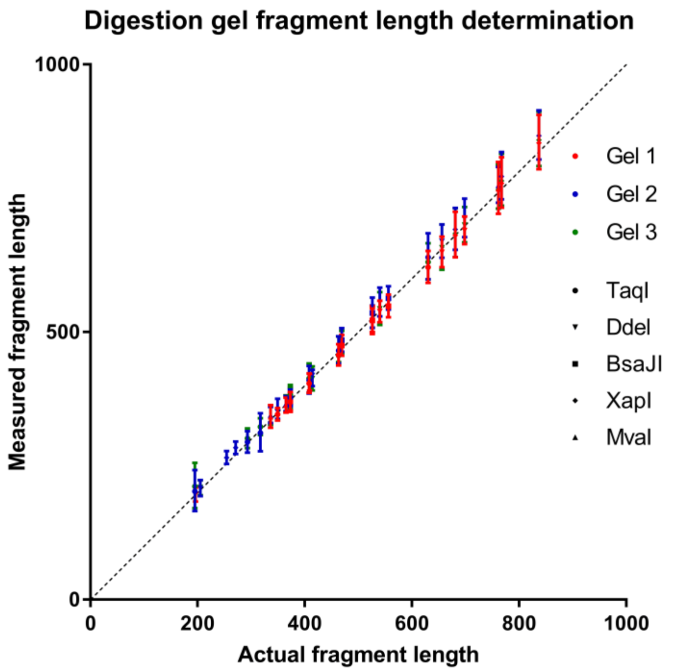


Supplementary Figure 13. **Determination of the resolving capacity of agarose gels with quantitative gel scanning.** Five restriction digests of pGL2-Control (Promega) were run in triplicate on 3 agarose gels. Individual measurements are plotted with standard deviations.

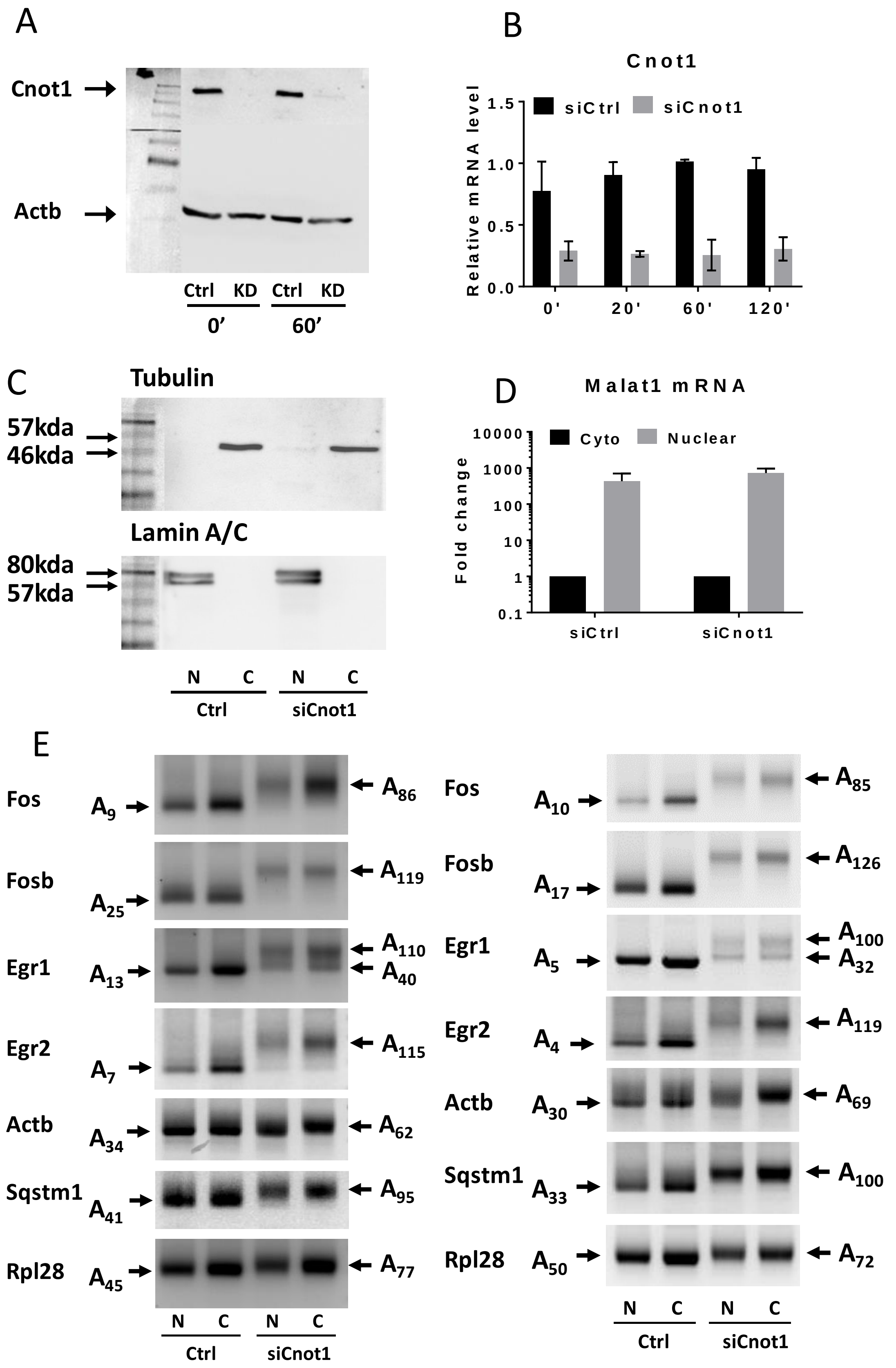


Supplementary Figure 14. **Controls and replicates for Figure 6.** Confirmation of knockdown of Cnot1 at the protein level by western (A) and at the mRNA level by RT-qPCR (B). Controls for the nuclear/cytoplasmic fractionation checked by western blot for tubulin (cytoplasmic) and lamin (nuclear) (C) and by RT-qPCR for Malat1 (nuclear non-coding RNA) (D). E. Two more independent replicates of the RL2-PAT for nuclear and cytoplasmic RNA of cells treated with control siRNA and Cnot1 siRNA.

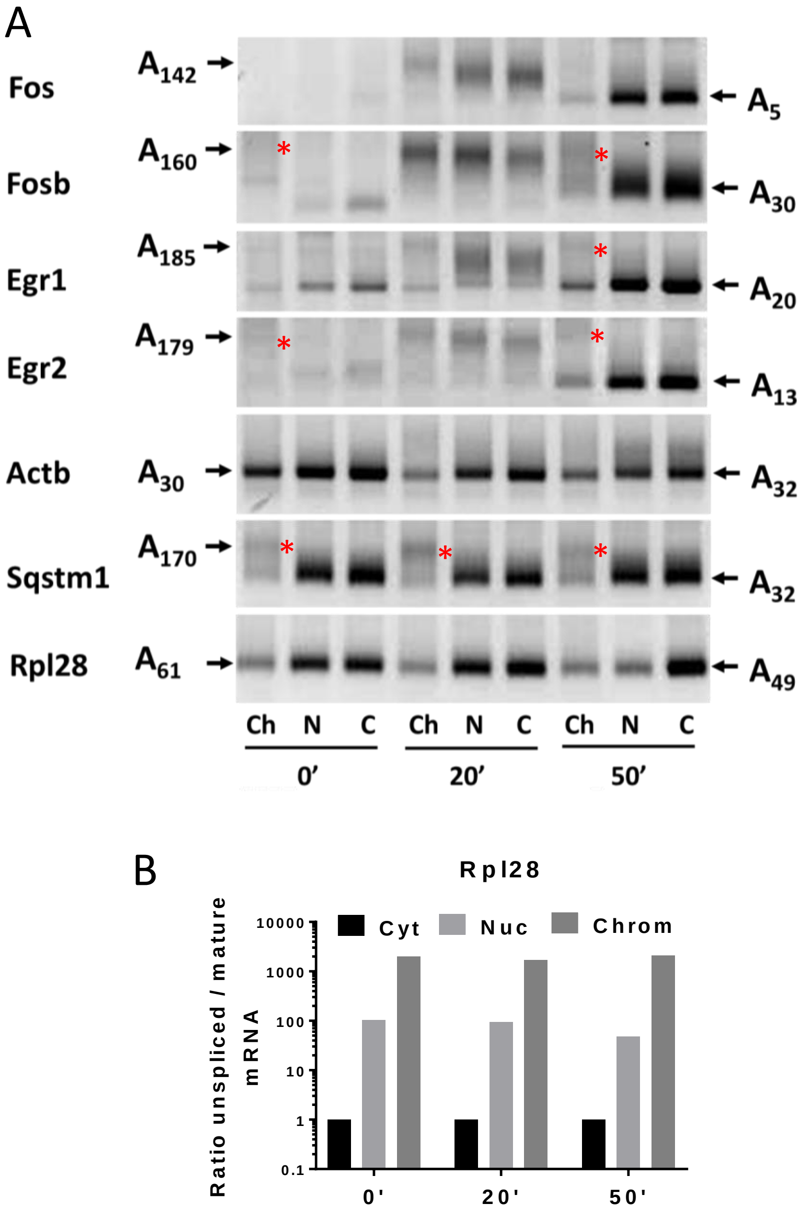


**Supplementary Figure 15. Replicate of Figure 7.** Most mRNAs with short poly(A) tails appear to have long-tailed precursors on the chromatin. A. Serum starved NIH3T3 cells were not treated (0’) or treated with serum for 20’ or 60’. Cells were fractionated into chromatin, nucleoplasm and cytoplasm and RNA was isolated. RL2-PAT was performed on this RNA for the indicated mRNAs. Red stars indicate potential precursor mRNAs with long tails. B Enrichment of uspliced Rpl28 mRNA in the chromatin and nucleoplasmic fractions..
