## Supplementary Tables for "Nuclear poly(A) tail size is regulated by Cnot1 during the serum response"

Supplementary Table 1.1

| | Immediate decay model best fit parameters | | | | | |
|---|---|---|---|---|---|---|
| | alpha | lambda | epsilon | RSS | Transcript | lifetime |
| Arc STC1 | 1.291023 | 0.035008 | 6.695178 | 7945.3278 | 5525.837 | 28.56508 |
| Arc STC2 | 1.207656 | 0.039539 | 6.695178 | 8635.1449 | 4132.223 | 25.29126 |
| Btg2 STC1 | 0.450648 | 0.049465 | 0.191004 | 7027764.4 | 3544.749 | 20.21625 |
| Btg2 STC2 | 0.588605 | 0.06568 | 0.191004 | 7850049.1 | 5246.837 | 15.2253 |
| Ccrn4l STC | 39.93553 | 0.020187 | 86.27389 | 246897.12 | 374040.5 | 49.53792 |
| Ccrn4l STC | 33.82768 | 0.018689 | 86.27389 | 570373.06 | 310707.6 | 53.50787 |
| Csrnp1 ST( | 4.957353 | 0.067725 | 32.6063 | 6284.622 | 23064.42 | 14.76549 |
| Csrnp1 ST( | 5.899285 | 0.086629 | 32.6063 | 2516.4594 | 22820.85 | 11.54345 |
| Ctgf STC1 | 0.677385 | 0.012761 | 99.99997 | 34127.931 | 71809.33 | 78.36162 |
| Ctgf STC2 | 0.43727 | 0.009608 | 99.99997 | 15653.275 | 49614.86 | 104.0766 |
| Cyr61 STC1 | 2.92945 | 0.045395 | 1.341972 | 297858215 | 165227.2 | 22.02877 |
| Cyr61 STC2 | 1.986544 | 0.039344 | 1.341972 | 195030933 | 118972.3 | 25.4169 |
| Dusp1 STC | 12.26347 | 0.110618 | 19.97353 | 19663384 | 1338433 | 9.040152 |
| Dusp1 STC | 12.1942 | 0.118817 | 19.97353 | 50476493 | 1680406 | 8.416305 |
| Dusp6 STC | 0.2731 | 0.04036 | 1.558522 | 12956.691 | 2804.539 | 24.77691 |
| Dusp6 STC | 0.192991 | 0.035295 | 1.558522 | 28632.653 | 2428.145 | 28.33272 |
| Egr1 STC1 | 9.813667 | 0.074994 | 57.01007 | 624857.05 | 407813.3 | 13.33448 |
| Egr1 STC2 | 8.08185 | 0.066611 | 57.01007 | 318911.52 | 326773.9 | 15.01252 |
| Egr2 STC1 | 1.226405 | 0.056423 | 0.2222 | 29859538 | 5513.549 | 17.72332 |
| Egr2 STC2 | 1.273512 | 0.056352 | 0.2222 | 13001435 | 5242.975 | 17.74568 |
| Ereg STC1 | 28.33946 | 0.005556 | 86.62286 | 394599.47 | 272265.1 | 180 |
| Ereg STC2 | 23.98453 | 0.005556 | 86.62286 | 394310.45 | 254071.2 | 180 |
| Fos STC1 | 2.786748 | 0.057162 | 12.17582 | 1422915.4 | 83302.44 | 17.49415 |
| Fos STC2 | 2.731225 | 0.066141 | 12.17582 | 1009602 | 84679.06 | 15.11922 |
| Fosb STC1 | 0.434245 | 0.027029 | 0.406141 | 4372420.9 | 2912.299 | 36.99762 |
| Fosb STC2 | 0.412392 | 0.028489 | 0.406141 | 2286283.6 | 2387.782 | 35.10184 |
| Gadd45g S | 0.243612 | 0.096474 | 0.502496 | 188640.89 | 2686.624 | 10.36547 |
| Gadd45g S | 0.126163 | 0.059156 | 0.502496 | 179604.92 | 1272.5 | 16.90446 |
| Hbegf STC | 10.98122 | 0.020424 | 50.74438 | 23093.273 | 48507.47 | 48.96095 |
| Hbegf STC | 14.48234 | 0.042198 | 50.74438 | 20116.35 | 51679.23 | 23.69789 |
| Id1 STC1 | 0.404496 | 0.052013 | 3.835974 | 25660.398 | 6673.699 | 19.22592 |
| Id1 STC2 | 0.359024 | 0.060429 | 3.835974 | 16666.046 | 7926.846 | 16.54825 |
| Id2 STC1 | 0.547999 | 0.029846 | 1.102898 | 271145.89 | 5617.094 | 33.50549 |
| Id2 STC2 | 0.583095 | 0.039514 | 1.102898 | 588926.91 | 7617.002 | 25.30748 |
| Id3 STC1 | 0.262372 | 0.045065 | 0.31131 | 426846.27 | 2028.452 | 22.19039 |
| Id3 STC2 | 0.266819 | 0.058987 | 0.31131 | 513415.72 | 2702.12 | 16.95296 |
| Klf9 STC1 | 1.132767 | 0.020435 | 99.99997 | 2694.2717 | 38583.78 | 48.93446 |
| Klf9 STC2 | 1.092296 | 0.018229 | 99.99997 | 5754.781 | 38824.75 | 54.85713 |
| Myc STC1 | 0.33852 | 0.054606 | 1.227544 | 264941.74 | 11931.56 | 18.31317 |
| Myc STC2 | 0.291501 | 0.047774 | 1.227544 | 425794.17 | 9493.379 | 20.93192 |
| Npas4 STC | 0.49522 | 0.016452 | 12.84932 | 3420.5901 | 2185.504 | 60.78251 |
| Npas4 STC | 0.372192 | 0.014533 | 12.84932 | 3590.6037 | 1950.45 | 68.80891 |
| Ptgs2 STC1 | 1.243045 | 0.017506 | 1.369972 | 134173532 | 123936.1 | 57.12403 |
| Ptgs2 STC2 | 1.731986 | 0.019929 | 1.369972 | 240604541 | 115264.9 | 50.17788 |
| Rgs16 STC | 0.814378 | 0.036062 | 6.433388 | 2418.4377 | 2217.03 | 27.72987 |
| Rgs16 STC | 0.343283 | 0.00871 | 6.433388 | 2596.1935 | 1186.365 | 114.8152 |
| Serpine1 S | 0.516 | 0.008119 | 0.682282 | 142388439 | 56288.92 | 123.1687 |
| Serpine1 S | 0.505808 | 0.008752 | 0.682282 | 295202088 | 50808.9 | 114.2648 |
| Trib1 STC1 | 1.524029 | 0.053825 | 10.64782 | 46150.278 | 28528.81 | 18.57875 |
| Trib1 STC2 | 1.299357 | 0.053923 | 10.64782 | 54683.056 | 26616.02 | 18.54486 |

Supplementary Table 1.2

| | Delayed decay model best fit parameters | | | | | | |
|---|---|---|---|---|---|---|---|
| | alpha | lambda | tau | epsilon | RSS | Transcripti | lifetime |
| Arc STC1 | 1.071444 | 0.036918 | 6.930377 | 6.695178 | 7868.1199 | 4586.074 | 34.01734 |
| Arc STC2 | 1.177338 | 0.04024 | 0.99519 | 6.695178 | 8639.1782 | 4028.494 | 25.84596 |
| Btg2 STC1 | 0.261711 | 0.249117 | 27.67087 | 0.191004 | 953760.43 | 2065.998 | 31.68506 |
| Btg2 STC2 | 0.291891 | 1.848072 | 28.22594 | 0.191004 | 5443509.1 | 2610.12 | 28.76704 |
| Ccrn4l STC | 39.59162 | 0.020121 | 0.327516 | 86.27389 | 246867.14 | 370819.7 | 50.02704 |
| Ccrn4l STC | 23.8529 | 11.55803 | 55.25475 | 86.27389 | 191006.16 | 219101.6 | 55.34127 |
| Csrnp1 ST( | 3.03744 | 0.10104 | 13.05546 | 32.6063 | 6191.6347 | 14137.31 | 22.95253 |
| Csrnp1 ST( | 2.842481 | 1.288798 | 21.42198 | 32.6063 | 1873.8482 | 11004.2 | 22.1979 |
| Ctgf STC1 | 0.539671 | 0.034123 | 40.76378 | 99.99997 | 27343.073 | 57350.69 | 70.06924 |
| Ctgf STC2 | 0.365666 | 0.038087 | 51.15572 | 99.99997 | 9659.9071 | 41613.42 | 77.41125 |
| Cyr61 STC: | 1.788504 | 0.088894 | 23.35512 | 1.341972 | 121274928 | 100933.9 | 34.60452 |
| Cyr61 STC: | 1.153555 | 0.090365 | 30.10443 | 1.341972 | 133924361 | 69156.48 | 41.17064 |
| Dusp1 STC | 6.166082 | 0.480805 | 16.20483 | 19.97353 | 8252397.4 | 672982.3 | 18.28468 |
| Dusp1 STC | 7.009078 | 0.209647 | 10.10748 | 19.97353 | 37676792 | 965856.6 | 14.87741 |
| Dusp6 STC | 0.195396 | 0.049732 | 14.21079 | 1.558522 | 11749.516 | 2022.884 | 34.31843 |
| Dusp6 STC | 0.1287 | 0.052641 | 22.0441 | 1.558522 | 24341.747 | 1643.174 | 41.04055 |
| Egr1 STC1 | 5.399557 | 0.176167 | 17.5154 | 57.01007 | 436001.11 | 224384.9 | 23.19183 |
| Egr1 STC2 | 4.849104 | 0.116123 | 15.38886 | 57.01007 | 252751.27 | 196064.6 | 24.00045 |
| Egr2 STC1 | 0.741493 | 0.095975 | 18.07403 | 0.2222 | 20808722 | 3336.299 | 28.49344 |
| Egr2 STC2 | 0.666864 | 0.553199 | 29.95134 | 0.2222 | 10674873 | 2748.809 | 31.75901 |
| Ereg STC1 | 23.63229 | 0.091373 | 102.7256 | 86.62286 | 204207.81 | 227051.5 | 113.6698 |
| Ereg STC2 | 19.75986 | 0.005582 | 498.5138 | 86.62286 | 242543.23 | 209327.8 | 677.6659 |
| Fos STC1 | 1.87799 | 0.081617 | 12.53039 | 12.17582 | 627537.73 | 56142.66 | 24.78268 |
| Fos STC2 | 1.666496 | 0.106448 | 14.33286 | 12.17582 | 683272.28 | 51674.67 | 23.72708 |
| Fosb STC1 | 0.297017 | 0.053814 | 30.10023 | 0.406141 | 1446070.3 | 1993.833 | 48.6826 |
| Fosb STC2 | 0.266607 | 0.058457 | 32.30647 | 0.406141 | 977486.22 | 1545.514 | 49.41301 |
| Gadd45g S | 0.144572 | 0.135438 | 9.852819 | 0.502496 | 195375.56 | 1604.129 | 17.23629 |
| Gadd45g S | 0.082512 | 0.069085 | 9.338697 | 0.502496 | 187296.19 | 840.6818 | 23.81364 |
| Hbegf STC | 9.296456 | 0.022779 | 11.93035 | 50.74438 | 22999.799 | 41069 | 55.82997 |
| Hbegf STC | 8.667402 | 0.728009 | 34.83298 | 50.74438 | 15397.265 | 30937.03 | 36.20659 |
| Id1 STC1 | 0.246944 | 0.095076 | 19.54376 | 3.835974 | 13697.949 | 4103.987 | 30.06162 |
| Id1 STC2 | 0.351985 | 0.06054 | 0.369018 | 3.835974 | 16671.43 | 7772.852 | 16.88691 |
| Id2 STC1 | 0.543634 | 0.029841 | 0.264209 | 1.102898 | 271155.76 | 5572.64 | 33.77558 |
| Id2 STC2 | 0.339197 | 0.11631 | 29.89068 | 1.102898 | 500536.65 | 4447.402 | 38.48842 |
| Id3 STC1 | 0.15334 | 0.10537 | 24.69535 | 0.31131 | 237226.09 | 1197.025 | 34.18576 |
| Id3 STC2 | 0.14482 | 0.15029 | 21.95593 | 0.31131 | 445114.69 | 1480.394 | 28.60972 |
| Klf9 STC1 | 0.811019 | 0.044625 | 33.93791 | 99.99997 | 1793.6885 | 27653.26 | 56.34689 |
| Klf9 STC2 | 0.826066 | 0.043481 | 36.66825 | 99.99997 | 4441.0184 | 29375.75 | 59.66666 |
| Myc STC1 | 0.26722 | 0.057444 | 5.812246 | 1.227544 | 262368.09 | 9435.904 | 23.22052 |
| Myc STC2 | 0.246316 | 0.048916 | 4.178563 | 1.227544 | 426314.95 | 8034.089 | 24.62157 |
| Npas4 STC | 0.418539 | 0.017709 | 13.73052 | 12.84932 | 3467.4986 | 1849.467 | 70.2003 |
| Npas4 STC | 0.291585 | 0.021791 | 28.99501 | 12.84932 | 3401.4195 | 1531.27 | 74.88643 |
| Ptgs2 STC: | 1.001012 | 0.021314 | 19.8984 | 1.369972 | 121053158 | 99880.54 | 66.81669 |
| Ptgs2 STC: | 1.287678 | 0.028608 | 26.86158 | 1.369972 | 199277370 | 85759.31 | 61.81624 |
| Rgs16 STC | 0.578865 | 0.43436 | 32.82203 | 6.433388 | 2266.2743 | 1588.113 | 35.12427 |
| Rgs16 STC | 0.31479 | 0.011171 | 18.97945 | 6.433388 | 2593.1203 | 1092.718 | 108.4945 |
| Serpine1 S | 0.502031 | 0.008138 | 3.577027 | 0.682282 | 142477494 | 54778.95 | 126.453 |
| Serpine1 S | 0.415976 | 0.051016 | 58.0937 | 0.682282 | 239083564 | 41850.38 | 77.69537 |
| Trib1 STC1 | 0.860776 | 0.281353 | 27.67168 | 10.64782 | 41194.894 | 16140.1 | 31.22593 |
| Trib1 STC2 | 0.650475 | 846.7088 | 34.11696 | 10.64782 | 45312.992 | 13361.69 | 34.11814 |

Supplementary Table 2. Parameters derived for the deadenylation models. Highlighted in yellow: Egr1 model timecourse 1 is an outlier for several parameters.

| | Egr1 | | Egr2 | | Fos | | Fosb | | Ptgs2 |
|---|---|---|---|---|---|---|---|---|---|
| | Time course 1 | Time course 2 | Time course 1 | Time course 2 | Time course 1 | Time course 2 | Time course 1 | Time course 2 | Time course 1 |
| Max decay rate $k^C$ ($min^{-1}$) | 1.30 | 0.72 | 0.95 | 1.31 | 1.08 | 0.77 | 1.18 | 2.90 | 0.33 |
| Max deadenylation rate, $k^A$, ($min^{-1}$) | 18.30 | 11.86 | 9.14 | 7.73 | 10.48 | 7.84 | 4.79 | 3.78 | 7.32 |
| Mean decay enzyme binding length, $\mu^C$ (bases) | 1.00 | 1.00 | 1.00 | 1.00 | 1.00 | 1.00 | 1.00 | 1.00 | 14.33 |
| Standard deviation of decay enzyme binding length, $\sigma^C$ (bases) | 40.30 | 18.26 | 16.51 | 11.11 | 23.56 | 18.34 | 15.06 | 13.45 | 26.73 |
| Mean deadenylation enzyme release length, $\mu^A$ (bases) | 82.17 | 0.03 | 12.60 | 0.00 | 0.00 | 0.00 | 4.94 | 13.68 | 0.00 |
| Standard deviation of deadenylation enzyme release length, $\sigma^A$ (bases) | 99.71 | 24.74 | 27.21 | 57.86 | 24.55 | 19.42 | 5.42 | 4.45 | 8.28 |
| Mean input length, $\mu$ (bases) | 183.55 | 185.18 | 201.71 | 187.61 | 142.65 | 155.88 | 173.29 | 153.79 | 220.47 |
| Standard deviation of input length distribution, $\sigma$ (bases) | 49.94 | 67.25 | 44.98 | 28.67 | 78.83 | 43.23 | 53.40 | 38.17 | 69.13 |

Supplementary Table 3. qPCR primer sequences

| Primer | Sequence | Amplicon size (bp) |
|---|---|---|
| Unspliced Fos F | TGACCGGAATGCTTCTCTCT | 165 |
| Unspliced Fos R | TGTCACCGTGGGGATAAAGT | |
| Spliced Fos F | GGGACAGCCTTTCCTACTACC | 87 |
| Spliced Fos R | GATCTGCGCAAAAGTCCTGT | |
| Unspliced Fosb F | GGGGTCGGTGTGTGTTATGT | 164 |
| Unspliced Fosb R | GATCCTGGCTGGTTGTGATT | |
| Spliced Fosb F | ACCCTCCGCCGAGTCTCAGT | 128 |
| Spliced Fosb R | TTGCGGTGACCGTTGGCACG | |
| Unspliced Egr1 F | GGGTCTCATCGTCCAGTGAT | 166 |
| Unspliced Egr1 R | GAAGCGGCCAGTATAGGTGA | |
| Spliced Egr1 F | AGTGATGAACGCAAGAGGCA | 121 |
| Spliced Egr1 R | TAGCCACTGGGGATGGGTAA | |
| Spliced Egr2 F | GTAGCGAGGGAGTTGGGTCT | 219 |
| Spliced Egr2 R | ATCATGCCATCTCCCGCCAC | |
| Unspliced Actb F | AAGATCTGGCACCACACCTT | 155 |
| Unspliced Actb R | TGAGAAGCTGGCCAAAGAGA | |
| Spliced Actb F | CTAAGGCCAACCGTGAAAAG | 104 |
| Spliced Actb R | ACCAGAGGCATACAGGGACA | |
| Unspliced Rpl28 F | CATCGTGTACACCTATTCCC | 88 |
| Unspliced Rpl28 R | ACGGTCTTGCGGTGAATTAG | |
| Spliced Rpl28 F | TACAGCACGGAGCCAAATAA | 74 |
| Spliced Rpl28 R | ACGGTCTTGCGGTGAATTAG | |
| Unspliced Sqstm1 F | GCCTCTGCTGCATTTTAGCCT | 137 |
| Unspliced Sqstm1 R | GAAAAGGCAACCAAGTCCCCA | |
| Spliced Sqstm1 F | CCTCTAGGCATTGAGGTTGA | 120 |
| Spliced Sqstm1 R | GCTTGGCTGAGTGTTACTCT | |
| Cnot1 F | AGCATAGCAGCATGTCTTCC | 147 |
| Cnot1 R | GCAGAACCCCCTGGTCTATC | |
| Malat1 F | GTGGGTGGGGGTGTTAGGTA | 176 |
| Malat1 R | CAACCTTCCTTAGCTGCCCG | |
| Gapdh F | AAGAAGGTGGTGAAGCAGGC | 114 |
| Gapdh R | ATCGAAGGTGGAAGAGTGGG | |

### Supplementary Table 4. Details of the NCounter codeset

| | | | | |
|---|---|---|---|---|
| | CODESET DETAILS | | | |
| | Customer Identifi | Accession | Position | Target Sequence |
| 1 | Agpat5_exon | Sin_Agpat5.1 | 1-100 | GACAAGCACGGGTCGCGCGGAGCAAAGCGAGCACCCCGAGGCGAGTGCGCCCGGCAAGCCGAGGCGTGCCCTTTCCAAGGCGGCGAGCAGAGGCCGTCAC |
| 2 | Agpat5_intron | Sin_Agpat5.1 | 895-994 | AGGAAATGAATACATTGGTTACAATAGGAGCCTCACTGTCGATACAGTTCTCTTCAGCTTGGACTGGGCTTCAATGTGGGCTGATCTCTTGTCAGATTGC |
| 3 | Arc_exon | Sin_Arc.1 | 138-237 | ATCACAGAACACGATCTGGCTTCCTCATTCTGCTCCAGTGTCCAGGGCTCTTTGGGTAATCAAGAAACCAAGTGTCTGAAAGGCAACAAAAAGTAGGCAC |
| 4 | Arc_intron | Sin_Arc.1 | 599-698 | GGTGTCATTGTAGTGGGCACTGGCACTTCACGGTCCAAGGACGGGGCTGAGGTGGGGGCGGTTCTGGGGGAATTTGCTAAGCCAGCTCACCGAGGACCCT |
| 5 | Arl6ip5_exon | Sin_Arl6ip5.1 | 1-100 | AAGGCGAAGCTCGCGAAGCAGAGGAACATGGACGTGAACCTCGCCCCGCTCCGTGCCTGGGACGATTTCTTCCCGGGCTCTGATCGTTTCGCCCGGCCGG |
| 6 | Arl6ip5_intron | Sin_Arl6ip5.1 | 305-404 | GCCTCAGGGAAGGGTCTCGGGAAGTGGCCCACCCAGTGGAAACTTGTTCTCAGGGCGAGGCCACAGCTCTCTGAGGTCCCCTTTCTTTCTCTTTTTGCTC |
| 7 | Atf3_exon | Sin_Atf3.1 | 1-100 | AAAATGATGCTTCAACATCCAGGCCAGGTCTCTGCCTCAGAAGTCAGTGCGACCGCCATTGTCCCCTGCCTCTCACCTCCTGGGTCACTGGTATTTGAGG |
| 8 | Atf3_intron | Sin_Atf3.1 | 756-855 | GCTCACAGCTATCTGTACCTTCAATTTCAGGAAATCCAACATCCTCTGCAGGCAACTGCATAAACATGTCACAAATACATAAATGCAGGTAAATACTCAT |
| 9 | Atox1_exon | Sin_Atox1c.1 | 29-128 | AGAAGGTCTGCATCGACTCTGAGCACAGCTCAGACACCCTGCTGGCAACCCTCAACAAAACAGGAAAGGCTGTTTCCTACCTTGGCCCCAAGTAGCCAGG |
| 10 | Atox1_intron | Sin_Atox1c.1 | 962-1061 | CTGACCCATGAATTACTGCAGTCACTTTGGTGGTTTGTGTTCTCTTTGGAACTGAGAAACACGAGGCCTGGTGAATGTGAGTGAGCTGACCAGAGCCCCA |
| 11 | Btg2_exon | Sin_Btg2.1 | 1-100 | GAGCCCGAGAGGTGGCCAGACCGTCATCATCGTTCTAATACAGCTACTTCCTCAGCCACCGGTATGAGCCACGGGAAGAGAACCGACATGCTCCCGGAGA |
| 12 | Btg2_intron | Sin_Btg2.1 | 604-703 | TCTTGAGTCCTAGCCACTGTTGCTGTATAGTCCTGAGCCTGTCATCATACAGGCCCATTCTGCCGCGTGTTTACCCGCTGCAAGTTGTAATCTGGGCCCC |
| 13 | Ccrn4l_exon | Sin_Ccrn4l.1 | 3-102 | CATGGGAAACGGCACCAGTCGACTCTACAGTGCTCTCGCCAAGACCGTCAACAGCAGTGCAGCTGCTCAGCACCCGGAGTACTTGGTGTCAACTGACCCC |
| 14 | Ccrn4l_intron | Sin_Ccrn4l.1 | 877-976 | TGCAGGTGAAGGGATGGTTCGGTGGTTGAGAGCAGGACTAACCTCCCACAGCAAACAGCTTCAGGGGATGCCCGGCCTCTGGCCTCTGTAGACACCTGCA |
| 15 | Cdc37_exon | Sin_Cdc37b.1 | 1-100 | CCCGGGTGGAACGCATGGAGCAGTTTCAGAAGGAGAAAGAGGAACTGGACCGGGGCTGCCGGGAGTGCAAGCGCAAGGTAGCCGAGTGCCAGCGCAAGCT |
| 16 | Cdc37_intron | Sin_Cdc37b.1 | 576-675 | GCACTTAGCGACGCCACCGCCTCCGATGTGGACAGGATGGGCGCGGCGGCCACGGCCCCCAAACCCCAGGCGCACCCCTGCCTTGCCCCAGCCCACTTGC |
| 17 | Cdkn2a_exon | Sin_Cdkn2a.1 | 1-100 | GGGACCCACTGGTCACACGACTGGGCGATTGGGCGGGCACTGAATCTCCGCGAGGAAAGCGAACTCGAGGAGAGCCATCTGGAGCAGCATGGAGTCCGCT |
| 18 | Cdkn2a_intron | Sin_Cdkn2a.1 | 694-793 | CACAGAATAAAAATAAACAAACGAGTTTGGGACCTTGAGATCATTTGCAAGTCAAGCGTGGAAACCTCTTTGATTCTAGACAGGTTGGGTGGTTGTGCGA |
| 19 | Csrnp1_exon | Sin_Csrnp1.1 | 132-231 | TCTCTCTCCTCCTTCCCCGCCTCTTCTGCCTCCCCTGCCTGGAACTCTGATGAGGAGGGACCAGGTGGTCAGGCACCCCAGTCTGATCAGGACTCCTGTG |
| 20 | Csrnp1_intron | Sin_Csrnp1.1 | 451-550 | CTGGCTCTGGGGCCTTAGAGGGGTGTTGCGGCTCTGTTTCCCCGGCACACCCTACCCCTTCCAGGAACACCACTGGGGCTCATCCCTCTGCTTCCACCCC |
| 21 | Ctgf_intron | Sin_Ctgfe.1 | 371-470 | TGACCCTAGCTCGTCACCTTGACATGTACAGTGATATAGCTAGCTGTTCTGATCCCTGTGACCCTACGCCTGACCTACAACTTTGTCTTCCTCTCCCGCA |
| 22 | Ctgf_exon | Sin_Ctgfe.1 | 43-142 | GCCGAAGCAGCGCCGCACTGCCCCGCCGGCGTGAGCCTGGTGCTGGACGGCTGCGGCTGCTGCCGCGTCTGCGCCAAGCAGCTGGGAGAACTGTGTACGG |
| 23 | Cyr61_exon | Sin_Cyr61.1 | 1-100 | GGGCAGACCGTGAGCGAGAGCGCCCCAGAGAAGCGCCTGCAATCTCTGCGCCTCCTCCGCCAGCACCTCGAGAGAAGGACACCCGCCGCCTCGGCCCTCG |
| 24 | Cyr61_intron | Sin_Cyr61.1 | 432-531 | CGGGACGCGATCGAGACACTTCTGGTGGACGGAGCCCCACAGCCAGACCTAGGCTCAGGGCAGTTTACGTGCCTCCCCGAGCAATGCAAAGCAATCTCAC |
| 25 | Dld_exon | Sin_Dld.1 | 6-105 | CCAATCGCGCTGCTCCTCGGCGATGACGTATATGTTGCGCAGGCGCAGGGTGGGGAGACCTTGGATAAGTCTTACCAGGCGTTCAGCGGAGGTGAAAACC |
| 26 | Dld_intron | Sin_Dld.1 | 824-923 | ACACGCAAAATGTCACTGTTTTAGGGAGAAGGTGAGCCGCGTCTCTCAGTACAGACAGTTCTTTGTTCCTCTTTGGTTGGCATATCTTGATCTCAGGTGC |
| 27 | Dusp1_exon | Sin_Dusp1.1 | 1-100 | GTGAAGGACTGGCGGGAGGAAAGCGCGGTGAAGCCAGATTAGGAGCAGCGAGCACTTGGGGACTTAGGGCCACAGGACACCGCACAAGATCGACCGACTT |
| 28 | Dusp1_intron | Sin_Dusp1.1 | 649-748 | TTAAGCCTTCCTTGTAAACTCGCAGCCCCTGGTGGGTGAGTGGCTTGTCTCCAGGTACTGTGTCGGTGGTGCTAATGAGAAAATATGTACCTTTGTATTC |
| 29 | Dusp6_exon | Sin_Dusp6.1 | 1-100 | ACGCTCAGACCCGTGCCCTTCGCGTCGGAAATGGCGATCTGCAAGACGGTGTCGTGGCTCAACGAGCAGCTGGAGCTGGGCAACGAACGGCTTCTGCTGA |
| 30 | Dusp6_intron | Sin_Dusp6.1 | 728-827 | TCTACCACTGCCGGGGGTTCCTAGGGTGCCTGTCCATAGTGTTCTTTGTGCTTCTCACTAGCTGTAAATTACTACAGCTTCTCCGGTTACAAGATTGCTT |
| 31 | Egr1_exon | Sin_Egr1.1 | 84-183 | GCCCCGGGCTGCGCCCACCACCCAACATCAGTTCTCCAGCTCGCTGGTCCGGGATGGCAGCGGCCAAGGCCGAGATGCAATTGATGTCTCCGCTGCAGAT |
| 32 | Egr1_intron | Sin_Egr1.1 | 1000-1099 | AACCACATCGGGAGTCAGTGGTAGCCGGGCGACCTCTTGCCAGGCCGCTTCGGGTCTCATCGTCCAGTGATTCCTCTCCAGTAACCAGGCCTCTCTGTTC |
| 33 | Egr2_exon | Sin_Egr2.1 | 1-100 | GATTAATAGCTGGGCGAGGGGACACACTGACTGTTATAATAACACTACACCAGCAACTCCTGGCTCCCCAACAGCCGGATCACAGGCAGGAGAGAGTCAG |
| 34 | Egr2_intron | Sin_Egr2.1 | 1035-1134 | CGGCGGAGCAGGGCAAGTAACAGCGAGAAGCGGCAAGGAGAATGCCTAGGGAGGCCCGCGCTGGCACCGGTGTGCGCCGGTTCCTTGGCGATGCTGTGTA |
| 35 | Ereg_exon | Sin_Eregc.1 | 33-132 | GTGCAGATTACAAAGTGTAGTTCTGACATGGACGGCTACTGCTTGCATGGCCAGTGCATCTACCTGGTGGACATGAGAGAGAAATTCTGCAGGTGAAGTG |
| 36 | Ereg_intron | Sin_Eregc.1 | 393-492 | AGTACCTCAGCAGCTACACTGATTGTCCTTTTGCTGCTATAGTTTGAACTAGATGCTGTGTAGAGCTAGTTTACTAATATTGAGTTAACTATGCATTCCA |
| 37 | Exosc9_exon | Sin_Exosc9c.1 | 1-100 | CTTATTGGTGGACCCCAATGAACGTGAAGAACGAGTAATGGATGGCTTGCTGGTGATTGCCATGAATAAGCATCGAGAAATTTGTACTATTCAGTCTAGT |
| 38 | Exosc9_intron | Sin_Exosc9c.1 | 361-460 | GTACCTCCCTACCCTGTGGGTGTACTAGCAGGACTAGGCACATCCTCTCCCGCTGAGACAAAACCAGGCAGTCCAAGTCAGGGCAGCACACCCAGTGGCA |
| 39 | Fos_exon | Sin_Fos.1 | 21-120 | AGAGCCGGCGGTTCCGCGAACGAGCAGTGACCGCGCTCCCACCCAGCTCTGCTCTGCAGCTCCCACCAGTGTCTACCCCTGGACCCCTTGCCGGGCTTTC |
| 40 | Fos_intron | Sin_Fos.1 | 423-522 | CAGGACTTGTCTGAGCGCGTGCACACTTGTCATAGTAAGACTTAGTGACCCCTTCCCGCGCGGCAGGTTTATTCTGAGTGGCCTGCCTGCATTCTTCTCT |
| 41 | Fosb_exon | Sin_Fosb.1 | 1-100 | TGGATCGCGCCCCTCTGCTCTCCCTCATTTCTTGGGACGCTTCGACAGGATTGGAACTGCATCTGAAGCGCACGCTGCACCGCGGCACTGCCCGGCGGGT |
| 42 | Fosb_intron | Sin_Fosb.1 | 935-1034 | ACCCTCCACCCAGCAAGAGTTAGGGCCTCGAACCCCGGTCACGCTGCCTCCGCCTCCTGCGCGGAGACGTAACGGGGGACCCGTGCGTAAAGGCTGACGC |
| 43 | Gadd45g_exon | Sin_Gadd45g.1 | 1-100 | CGCACTCGCTGGTGGTGGGCGCGCATCGGACTCTGGGAATCTTTACCTGCGCTCGGGTTCCCTCCGCACTCTTTTGGATAACTTGCTGTTCGTGGATCGC |
| 44 | Gadd45g_intron | Sin_Gadd45g.1 | 411-510 | TGGGTGATGGGGAATGAGGGCGCCAGATCATGACTAGGGTGCCAGAGGTGTGGGCGTCAGGTCGGGACTGGGGATCCGGGTGTGTGGGCGCCAGGTCGGG |
| 45 | Grrp1_exon | Sin_Grrp1.1 | 540-639 | GACGCACCAGGTGATTGTACGTCGTCAGGAGCCAGCCCTGCGCGGGGGACCAGGACCGCTGCACCACACCCCTGCAATGAGTTGGGGGCCTCTGCATCGC |
| 46 | Grrp1_intron | Sin_Grrp1.1 | 7-106 | CTTGGGTTTGAAGCTACACTGTTGGGAAGAGGTATTTTTTCCGGTTAAGGCCGGTTTCGGGAACCGGGCTAGGACCCCGCCTGATTTTCCTTTCCAAGTG |
| 47 | Has2_exon | Sin_Has2.1 | 142-241 | CCAGCTGTTGACCCTTATGACATGCCAGGAACCAGCTACTCAACCCCAGGCCTGAGTGCCTACAGCACTGGCGGGGCAAGCGGAAGTGGTGGGCCTTCAA |
| 48 | Has2_intron | Sin_Has2b.1 | 879-978 | TACTTTTCCACTGGGCATCTTCCCGAGCGTCAAAGGGGCTCCGGAGTCAGAAAACAAGGATTTCCACTGTGTACGGAAAATTCATCCCACAACTGTCCTT |
| 49 | Hbegf_exon | Sin_Hbegf.1 | 2-101 | TTATTCCCGGCCGCGCGCTGCGAGCGAGAGCTCAGTCGGCCCGGGGAGCTGCACGCGGCTGGCTGGTCGGCCTGACAGACCTTCAAGGGCTGGAGTGGAC |
| 50 | Hbegf_intron | Sin_Hbegf.1 | 440-539 | ACTTGGTGGGGCCCGTGCGCCCTGAAGGGCGGGCGTGGACCTCGATGGCGTGGTAGAGAGGGGTCTAAGCAAAGGGTCTTATTGCGGCCTCCTCTCTTCT |
| 51 | Id1_exon | Sin_Id1.1 | 22-121 | GTGCCGCAGCCGCTGCAGGCCCTAGCTGTTCGCTGAAGGCGGGCAGGACAGCGGGCGAGGTGGTACTTGGTCTGTCGGAGCAAAGCGTGGCCATCTCGCG |
| 52 | Id1_intron | Sin_Id1.1 | 528-627 | AGCTCGCATTTTCATCGTGCCTCCTGGAGTAGAGAAATGGGAACGCCTCTCCCCTCCTTGTCCTTTCCAGTGGGTCTCATCCCTTATCTCGCTCTGGTGT |
| 53 | Id2_exon | Sin_Id2.1 | 38-137 | TGTCGGACCACAGCTTGGGCATCTCCCGGAGCAAAACCCCGGTGGACGACCCGATGAGTCTGCTCTACAACATGAACGACTGCTACTCCAAGCTCAAGGA |
| 54 | Id2_intron | Sin_Id2.1 | 513-612 | CTGCGCTGTAGCATGATCCTGTTTCATGGGACTTGTTGGCACTTTTGTGAAAGGAGGAGGGGGGCTACCTTCCTTTAAGATTACCTAATATCCTGCCTTT |
| 55 | Id3_exon | Sin_Id3.1 | 1-100 | TAGCCTGGCCATTGCGCGAGGCCGCGGTAAGAGCCCGTCGACCGAGGAGCCTCTTAGCCTCTTGGACGACATGAACCACTGCTACTCGCGCCTGCGGGAA |
| 56 | Id3_intron | Sin_Id3c.1 | 57-156 | GAGTTATCAGCTGGAGGTAGAGACCGAGTTTCCTCCCTGGCGCCGGCCAGTCTGCGGCATCCTCCGCCTGGGCGCGCTCGGCGGAAACTGACGGCTCCCT |
| 57 | Il6_exon | Sin_Il6.1 | 27-126 | ATGCTGGTGACAACCACGGCCTTCCCTACTTCACAAGTCCGGAGAGGAGACTTCACAGAGGATACCACTCCCAACAGACCTGTCTATACCACTTCACAAG |
| 58 | Il6_intron | Sin_Il6.1 | 527-626 | GGACCAGAAAGTGCTTTTTGGCTGAATGTAGTAAGTTCTGTGTGTGTGAAATCAGGATGCTCTAGGGTCAGCCCAGATAAGATGCATATTGTATAGAGCC |
| 59 | Itga1_exon | Sin_Itga1c.1 | 16-115 | GAAGCTCGGGGGGCCAGAAGGGGAGTGAAAAAAGTCATGGTTATTGTGACTGATGGAGAATCGCATGACAACTATCGCCTGAAACAGGTCATCCAAGACT |
| 60 | Itga1_itnron | Sin_Itga1c.1 | 286-385 | TCGTGGTATTTCTGTAGCCCTTCTATCTGTCTGCTTACGCTGTCCCAGGTTGTCACATTTCCCAGGTGTAAAGGGAATGGAAGGCAGGGTGTAGGAAGGG |
| 61 | Klf9_exon | Sin_Klf9.1 | 44-143 | CGTGCCGGAGCACGGGGGCGCTCCGGAAGCCGAGCGGCTGCGACTACCTGAGCGCGAGGTGACCAAGGAACACGGTGACCCGGGGGACACCTGGAAGGAT |
| 62 | Klf9_intron | Sin_Klf9.1 | 896-995 | TCGTTTTAGGATGACTAAGGACTCATTTAACTGGATGGCGGCTACTTTTCAGGGCTTGCCAAGGTGGTTTTAAATTTGGCTAAGAGTCCCCTGAGAGGTT |
| 63 | Mapk8_exon | Sin_Mapk8b.1 | 27-126 | AGACTAATTGCTTGCCATCATGAGCAGAAGCAAACGTGACAACAATTTTTATAGTGTAGAGATTGGAGATTCTACATTCACAGTCCTAAAACGATACCAG |
| 64 | Mapk8_intron | Sin_Mapk8b.1 | 689-788 | CATATACTGATATCAGGGAAGAGGAAATGGCAGGCTTGTTCGTCCATGGATGCATCGTTTCCCATAGCTGATTAGTGAGATGTCTCATGGAATCTTCATT |
| 65 | Myc_exon | Sin_Myc.1 | 1-100 | CGCTGTAGTAATTCCAGCGAGAGACAGAGGGAGTGAGCGGACGGTTGGAAGAGCCGTGTGTGCAGAGCCGCGCTCCGGGGCGACCTAAGAAGGCAGCTCT |
| 66 | Myc_intron | Sin_Myc.1 | 676-775 | ATTGATATGTGTCCTTTGAGGGGTCAAACCGGGAGGTCGCTTCGTGGTGGCCAAAGAAAGCCCTTGGAATCCTGAGGTCTTTGGAGAAGGGATTACCTTT |
| 67 | Myl12a_exon | Sin_Myl12j.1 | 728-827 | TTTGGAAGATAGTTCCTGCAGAATTGTGACCCTGCATTAGCTCTGCCGCCAGCTGGACGGACCTGCTTTACCTCGTCCATGTAAACTTCCTGCCACCGAA |
| 68 | Myl12a_intron | Sin_Myl12a.1 | 248-347 | TTCATGAAACAGTGTCTCCTCCTGGCACACGAGTATTCCTGTGCCTGTCTCTGGATTGGACTGGAGGGCTTGTGTATCTGCGTTCCTCCTCCATGGATGT |
| 69 | Npas4_exon | Sin_Npas4.1 | 21-120 | CGGAGCCCAGGAGTGGAGCGAGAGCGAGCAAGAGCCTGAGCGAAAAGACCGGGAAGCAAGGAAGAGGAAGCCTCCGGTGCATCGGGAAAGGATCGCAGGT |
| 70 | Npas4_intron | Sin_Npas4.1 | 664-763 | AGGAAGAGTAGCTTGTGCTAGAAAATGCTGAGAGGGTCTTTCTATGCGCTCTGGAGCTGTTCGACGTCCTGAAGCCATCACCCTTTCTGGCGCTGCCCGC |
| 71 | Nppb_exon | Sin_Nppb.1 | 27-126 | ACCCCTACTCCGTGAAAAGGTCTGGCCGGACACTCAGCCCCAGTATAAAAGGCAGAGGCACCGTTGTTGAAGACACCAGTGCACAAGCTGCTTGGGGAGG |
| 72 | Nppb_intron- 2 b | Sin_Nppb.1 | 398-497 | AAGATTGGGCAGCATATCTCTTAACTGATGAGCACTATGGAAGGATGGGGGATTCAGGTGTGTGTGTTTCTGACGTCTGGGCTCCCCAATCCATCACAGA |
| 73 | Per1_exon | Sin_Per1b.1 | 1-100 | TTCTGGGTTGCGGGCCGAAACGGCAAGCGGATGGAGGGCGCTCGAACGGCCAGGTAGGGATCCCTGCTGCAATCCTAGGGTTAGTCTTTGTCCCGGAGCT |
| 74 | Per1_intron | Sin_Per1b.1 | 950-1049 | GTGTGTTGGCCAGTATAGGACTGGGTCTGTTTCCCACTTCCACGAAGATGGGGATTGGGGGAGGAGTCGTTCCTGCCCTCCTGTGGTCCCTCCAGCAACC |
| 75 | Plk2_exon | Sin_Plk2b.1 | 1-100 | CGCACAAGCGAAGCAGGACGTCAGACTAGAGAGTAGGGAGAGAGACTGGTGCTCGAGGGACAGGGCTAGCCCGGACGCTTGTCCGCGCCTCGGAGGTGGC |
| 76 | Plk2_intron | Sin_Plk2b.1 | 780-879 | AATGTGCTGCCCTTCCTGAAAAACCATTATACAAAGCAGATTGTACTGCTCTGCCCACCCTCTGTTCCTCAAATTTTGCTTCCCGTTCCCATTTACGTAT |
| 77 | Ptgs2_exon | Sin_Ptgs2e.1 | 442-541 | TCATCGGTGGCATTGGGAAGTGAACCTTAGAAGTTTGTGCTTGCCAGGCACTGGTCTCTGCTATTGAGGCAGAAGAGGTCCAGCCTTTACACACACACAC |
| 78 | Ptgs2_intron | Sin_Ptgs2e.1 | 66-165 | CCCCAAGGCTCAAATATGATGTTTGCATTCTTTGCCCAGCACTTCACCCATCAGTTTTTCAAGACAGATCATAAGCGAGGACCTGGGTTCACCCGAGGAC |
| 79 | Rgs16_exon | Sin_Rgs16.1 | 1-100 | CTCCAGACCTCTTCTGCGCAGCCTGGTACTTGCTACTCGCTTTTCCCTCTTCGCCCTCCGGTCCGCCACCATCCTCTCCACGACGTGCTGTCCTGCGTCC |
| 80 | Rgs16_intron | Sin_Rgs16.1 | 397-496 | GACTGAACTCTGGGTGGTGGTGTTAGTTGGGAGGCCCTCCTTCCCCAACTTCTCGGTGCTGGCAAACAAACAGAATCCTTACACCTTGATGGTTTTTCTT |
| 81 | Serpine1_intron | Sin_Serpine1.1 | 478-577 | GAAATCAGGGATCAACTTTCAGGAATCAGCTCTCACTTTAAGCCAAAAGCTATCATCGTCTAAAGGTCTCTTACCCCACAGAAGGGATCAGAGATGCACA |
| 82 | Serpine1_exon | Sin_Serpine1.1 | 10-109 | CACAGCTGGATCAGGCTGCAGCAGAGCCCCGAGAGCTTTGTGAAGGAGGACCGCCGCACACCCGCCTCCGGCAGCACACAGCCAACCACAGCTGAGCGAC |
| 83 | Sgk1_exon | Sin_Sgk1.1 | 67-166 | TGGGGAGTCAGACTTTGAGGCCATGTGTCAATCATGCCTGGGTGACCATGCTTTCCAAAGGGGGATGCTCCCTCCAGAGGAGTCCTGTTCCTGGGAGATC |
| 84 | Sgk1_intron | Sin_Sgk1.1 | 664-763 | GAGGGTGACTTGCATAAGTTTCACTCTCTCCTTTATTATGTGGGTCCTGAGGACTCAGGTCACTGGGCCTTAACCCACAAAGCCATCTCACCAGCCTCCT |
| 85 | Tapbp_exon | Sin_Tapbp.1 | 2-101 | TAGCGACCGTCGTCTCCGTCGTCTCGGCTGGACCAGAGGCGATCGAGTGCTGGTTCGTGGAGGATGCAGGTGGGGGTGGCCTGTCTAAGAAACCTGCCAC |
| 86 | Tapbp_intron | Sin_Tapbp.1 | 269-368 | GGATGCTCTCTAAGTCGTTCACTACTTTGCTAGTTTGTGCCTTCCGGCCTGTGTGTAGTCAGCACATTTCCAGTCCGGGCTACCGTTACACCCTTTTCGG |
| 87 | Trib1_exon | Sin_Trib1.1 | 8-107 | GGCGCCTCGCAGCCCCGCGGTCCGGGTCTACTGTTTCCGGCTGCCCGGGGAACCCCGGCCAAACGCCTGCTGGACACGGACGACGCGGGGGCAGTGGCGG |
| 88 | Trib1_intron | Sin_Trib1.1 | 702-801 | GACGCCCTGCAGTGTGTGCCTCCTCGGGACTTGGAGCAACTGGCAAGCTCTTTGCGCGAGGCTCGCGGCATGGGTTCTGTATAGACCAAGTCGGGCTTCT |
| 89 | Uchl5_exon | Sin_Uchl5.1 | 1-100 | GGGGTTGACGCTCGGCGGCGGGTTTTGCTGAGGTCTGCGGCCGGTGGCAGCTTGTGCTGCGGGGCTCCGGAGAGCGCGAGGTGGAAGCGGGGCGGTGTGT |
| 90 | Uchl5_intron | Sin_Uchl5.1 | 486-585 | TCTCCCTCCCTCATCATCTCCTCCGCGCTTCTCTTGTCCTTTCCACTGTCCCTTCCCTGCGGAGCCTCCCTGTCTCCAGTGTGGTCAGGTGCATCGCTCA |

Table 5. RL2-PAT primer sequences

| Supplementary Table 5. Sequences of PAT primers | | |
|---|---|---|
| **Name** | **Sequence (5'→3')** | **Expected size of deadenylated product** |
| Fos 3F1 | CAGCGTCAATGTTCATTGTCA | 280 |
| Fos 3F2 ('Fos PAT') | CTGACATTAACAGTTTTCCATG | 221 |
| Fosb | ATTGACTCCATAGCCCTCAC | 184 |
| Egr1 | AGCTGAGCTTTCGGTCTCCA | 314 |
| Egr2 | GTGCTTCAATGTCACTGCCG | 187 |
| Id1 | CAGCCTCCAGAGACTTTGGG | 246 |
| Ptgs2 | CCTGCTGATTGAACCTGGGA | 205 |
| Actinb1 3f1 | AAACTTTCCGCCTTAATACTTC | 219 |
| Actinb1 3f2 | GGAGGATGGTCGCGTCCAT | 308 |
| Actg1 3f1 | ACCACCATCGGTTGTTAGTTG | 222 |
| Actg1 3f2 | GTTGGGAACGTTGCATCGA | 195 |
| Sqstm1 3f1 | AAGAGGGGACTGTCCATAGT | 255 |
| Sqstm1 3f2 | TTGCACCAGCAGTCCAGAAT | 358 |
| Rpl28 3f1 | GCCACTTCTTATGTGAGGAC | 254 |
| Rpl28 3f2 | TCGTGGTAGTTATGAAACGCA | 295 |
| Rps4x of1 | TTATTGGCAAGGGTAACAAAC | 221 |
| Rps4x of2 | CCGGCTCTTTTGATGTGGTT | 296 |
| PAT Anchor sequence | 5'-rApp GGT CAC CTT GAT CTG AAG ddC- 3' | |
| PAT reverse primer (PATR1) | 5'-GCT TCA GAT CAA GGT GAC CTT TTT- 3' | |
